# Meaning-based guidance of attention in rhesus monkeys during naturalistic scene viewing

**DOI:** 10.64898/2026.03.11.711223

**Authors:** Orhan Soyuhos, Taylor R. Hayes, Wenqing Hu, Taylor P. Hamel, Brinda Sevak, John M. Henderson, Xiaomo Chen

**Affiliations:** Center for Neuroscience, University of California, Davis, Davis, CA, USA; Department of Psychology, University of California, Davis, Davis, CA, USA; Center for Mind and Brain, University of California, Davis, Davis, CA, USA; Department of Neurobiology, Physiology and Behavior, University of California, Davis, Davis, CA, USA; Department of Biomedical Engineering, University of California, Davis, Davis, CA, USA

## Abstract

High-acuity vision in humans and other primates is restricted to the fovea and requires frequent saccadic eye movements to sample visual information, a process known as overt attention. Classical visual salience theory explains how low-level image features guide these movements, but recent human studies have shown that overt attention is also strongly guided by scene meaning—the spatial distribution of local scene informativeness as judged by human observers. It remains unknown whether this form of attentional guidance is uniquely human or extends to non-human primates. We recorded eye movements from two rhesus macaques freely viewing naturalistic indoor scenes. Fixation selection was modeled using meaning maps alongside image-based salience maps and center proximity. In both monkeys, meaning robustly guided fixation selection after controlling for visual salience and center bias. Moreover, attention was directed to high-meaning regions even when their visual salience was low, whereas salience played an increasingly important role as meaning decreased. While this prioritization of meaningful regions remained robust across environments, long-term familiarity with the photographed environment reduced the influence of meaning on fixation selection. Finally, the influence of meaning increased with attentional engagement, robustly in one monkey and as a trend in the other. These findings suggest that macaques share the human attentional bias toward informative scene regions and establish a behavioral foundation for investigating its neural mechanisms.

## Introduction

Real-world visual scenes contain far more detail than can be seen clearly at any given moment (Henderson, 2003). High-acuity vision in humans and other primates is limited to the fovea. They therefore sample the visual environment with frequent saccadic eye movements, a process known as overt attention (Findlay & Gilchrist, 2003; Henderson, 2013). Classical models of attentional selection emphasize the role of visual salience in guiding attention during naturalistic scene viewing, proposing that gaze is drawn toward locations with prominent local contrast in visual features such as luminance, color, and orientation, which are combined into a salience map that guides stimulus-driven attention (Itti & Koch, 2001; Parkhurst et al., 2002; Treisman & Gelade, 1980; Wolfe, 1994). Although these models successfully explain salience-driven aspects of fixation behavior, they do not fully account for gaze patterns during free-viewing, in which both visual salience and meaning contribute to attentional selection.

A major obstacle to evaluating the role of meaning in attentional guidance has been the lack of methods for quantifying it in visual scenes (Henderson, 2017). The meaning map approach was first developed to systematically quantify the spatial distribution of local informativeness across real-world scenes (Henderson & Hayes, 2017; Henderson et al., 2019). In this approach, human raters judge how informative or recognizable small patches of a scene are when the patches are viewed out of context, and the ratings are reassembled into a map that grades every location in the scene by its informativeness, which we refer to as scene meaning. However, this method relied on labor-intensive human rating procedures. More recently, the DeepMeaning model provided an image-computable extension of this framework, trained to reproduce human meaning maps from the features of a pretrained vision–language model (Hayes & Henderson, 2025). Using these approaches, scene meaning has been shown to predict fixation selection beyond low-level salience in human adults (Hayes & Henderson, 2022a, 2022b; Henderson & Hayes, 2017, 2018; Zhang et al., 2022) and infants (Oakes et al., 2024), even when meaning is task-irrelevant (Peacock et al., 2019). Notably, meaning-based guidance emerges rapidly, influencing the earliest fixations (Henderson & Hayes, 2017, 2018). Consistent with this early influence, neural signatures associated with the processing of scene meaning appear shortly after visual onset in EEG recordings (Kiat et al., 2022). Together, these findings suggest that attentional selection during naturalistic viewing is robustly shaped by high-level scene meaning in humans.

Whether meaning-based guidance of attention reflects a uniquely human strategy or a conserved feature of the primate visual system remains unknown. Non-human primates provide a critical model for investigating the circuit mechanisms underlying visual attention, given their close anatomical, functional, and behavioral similarities to humans (Bichot et al., 2015; Buschman & Kastner, 2015; Buschman & Miller, 2007). Extensive work in macaques has identified attentional control signals distributed across interconnected cortical and subcortical networks, including frontoparietal, temporal, and midbrain structures, that support both visual salience-driven and goal-directed attention (Boshra & Kastner, 2022; Moore & Zirnsak, 2017; Xia et al., 2024). In parallel, neurons throughout the visual hierarchy exhibit selectivity for complex object features and higher-order visual content, indicating that primate visual representations extend well beyond low-level image statistics (Peelen & Kastner, 2014; Ponce et al., 2019; Taubert et al., 2017; Vinken et al., 2025). In natural-scene search, macaque saccades are guided by scene context and by expected reward rather than by image features alone (Segraves, 2023; Segraves et al., 2017). However, these studies leave open whether scene meaning itself guides attention in non-human primates when they view natural scenes freely, without a search task.

Here, we examined the extent to which overt attention in macaques, measured as fixation selection, is guided by scene meaning and whether this guidance depends on behavioral context. We recorded eye movements from two rhesus macaques freely viewing naturalistic indoor scenes and modeled fixation selection using meaning maps, low-level salience maps, and spatial center bias. To assess contextual modulation of meaning-based guidance, we compared gaze behavior during viewing of unfamiliar and familiar scenes and across different levels of attentional engagement, quantified as the total time spent fixating the scene during a trial. We found that scene meaning robustly guided fixation selection in macaques and exerted an influence comparable to that of low-level salience after accounting for center bias. As in humans, this guidance was present from the earliest fixations of a trial. In addition, meaning interacted with visual salience in guiding attention, such that high-meaning regions drew fixations even when their salience was low, whereas salience played an increasing role as meaning decreased. Moreover, attentional guidance was modulated by behavioral context: meaning guided fixation in both unfamiliar and familiar scenes but more strongly in unfamiliar ones, with familiarity increasing the likelihood of fixations on low-meaning areas. Finally, the influence of meaning increased with attentional engagement, robustly in one monkey and as a trend in the other. Overall, these findings suggest that macaques share the human attentional bias toward informative scene regions and establish a behavioral foundation for future mechanistic investigations into how scene meaning is represented and utilized by neural circuits.

## Results

We recorded eye movements from two male rhesus macaques (Monkey V and Monkey I) freely viewing naturalistic indoor scenes. The stimulus set consisted of 200 images evenly split between unfamiliar and familiar scenes. Unfamiliar scenes depicted indoor environments the monkeys had never experienced, such as restaurants and residential interiors, whereas familiar scenes were photographs of their housing and laboratory surroundings. Each trial began with central fixation followed by 5 s of free-viewing (Figure 1A). In total, we analyzed 3140 fixations from Monkey V and 2932 fixations from Monkey I across 193 and 175 images, respectively, all from the first viewing of each scene (Figure S1).

**Figure 1.**
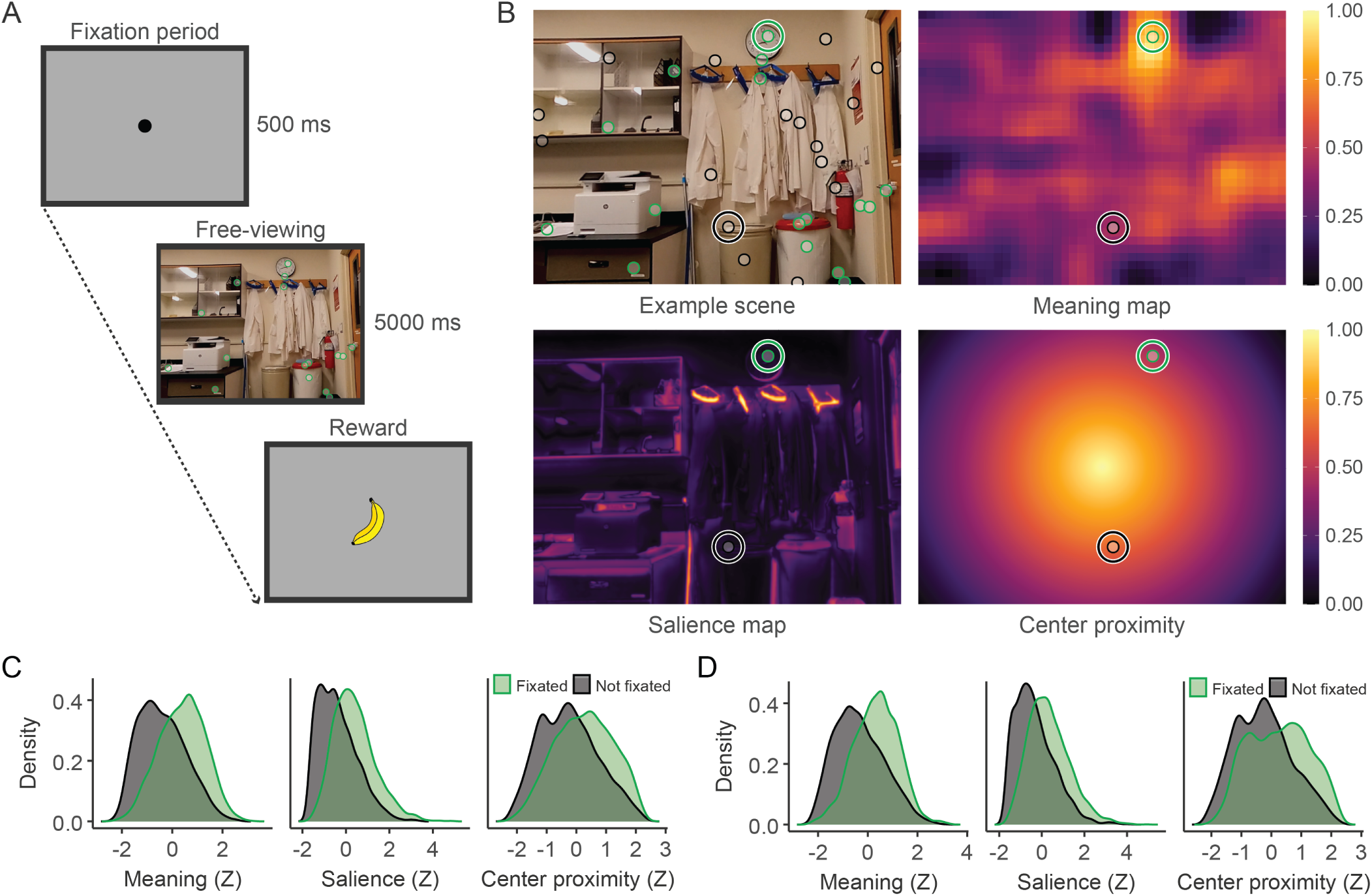
Experimental task and spatial predictors of fixation selection. (A) Behavioral paradigm. Monkeys initiated each trial by fixating a central point (500 ms) before freely viewing a naturalistic scene for 5 s to receive a juice reward. (B) Example scene and corresponding spatial predictor maps. The top-left panel shows an example scene with fixation locations (green dots) and randomly sampled non-fixated regions (black dots). Together, these locations provide an account of regions that the subject did and did not fixate. Each location was used to compute a mean value from the meaning, salience, and center proximity maps across an approximately 3-degree window, illustrated by the circles around example fixated (green) and non-fixated (black) locations. These spatial maps respectively visualize the spatial distribution of local scene informativeness, local contrast in low-level features, and the central fixation bias used to model fixation selection. (C) Predictor distributions for Monkey V. Density plots comparing the normalized (*z*-scored) values of meaning, salience, and center proximity at actual fixated locations (green) versus randomly sampled non-fixated locations (black). (D) Predictor distributions for Monkey I. Same as in (C).

### Scene meaning guides overt attention during free-viewing

To determine whether overt attention in macaques is guided by scene meaning, we quantified fixation selection as a function of three spatial predictors and their interactions: local meaning, image salience, and center proximity (Figure 1B). Local meaning quantified the informativeness of each location, as estimated by the meaning map of each scene (Hayes & Henderson, 2025; Henderson & Hayes, 2017). Image salience captured local contrast in low-level features such as luminance and edge orientation (Itti et al., 1998). Center proximity measured closeness to the image center to control explicitly for the known bias toward central fixation, independent of image content (Hayes & Henderson, 2022b; Tatler, 2007). The three predictors had low variance inflation factors in both animals (maximum VIF = 1.33; Figure S2A).

For each predictor, values at fixated locations were compared against those at randomly sampled non-fixated locations (see Methods; Figure 1C and D). If scene meaning guides attention in macaques, then locations with higher meaning values should be more likely to be fixated, even after accounting for salience and center bias. To test this, we fitted a Bayesian Bernoulli generalized linear mixed model (GLMM) predicting whether a sampled location was fixated based on meaning, salience, and center proximity, with a random intercept for scene (Table 1; Figure 2A and B; Fixation model). For Monkey V, meaning showed a robust positive effect (*β* = 0.73, 95% HDI [0.64, 0.83], *P* (*β >* 0) *>* 0.999), after controlling for the positive effects of salience (*β* = 0.71, 95% HDI [0.64, 0.79], *P* (*β >* 0) *>* 0.999) and center proximity (*β* = 0.30, 95% HDI [0.23, 0.36], *P* (*β >* 0) *>* 0.999). Monkey I showed a similar pattern, with meaning acting as a strong predictor of fixation (*β* = 0.46, 95% HDI [0.38, 0.54], *P* (*β >* 0) *>* 0.999) alongside salience (*β* = 0.58, 95% HDI [0.51, 0.66], *P* (*β >* 0) *>* 0.999) and center proximity (*β* = 0.39, 95% HDI [0.32, 0.45], *P* (*β >* 0) *>* 0.999). Notably, meaning remained a credible positive predictor in both animals when locations were sampled at smaller spatial scales (Figure S3). In a model-free analysis, the fixation density of each scene also correlated with its meaning map (*r* = 0.27 and 0.24), even after the salience and center proximity maps were regressed out of it (semipartial *r* = 0.14 and 0.13, both *p <* 0.001; Figure S4). To interpret the magnitude of these effects, we converted the model coefficients into predicted fixation probabilities. For a region with average features, the baseline probability of fixation was approximately 51% for both animals. Increasing meaning by 1 standard deviation (SD) raised this probability to 68% in Monkey V and 63% in Monkey I. In comparison, increasing salience by 1 SD yielded probabilities of 68% in Monkey V and 65% in Monkey I (Figure 2C and D). Thus, meaning exerted an influence comparable to that of visual salience.

**Figure 2.**
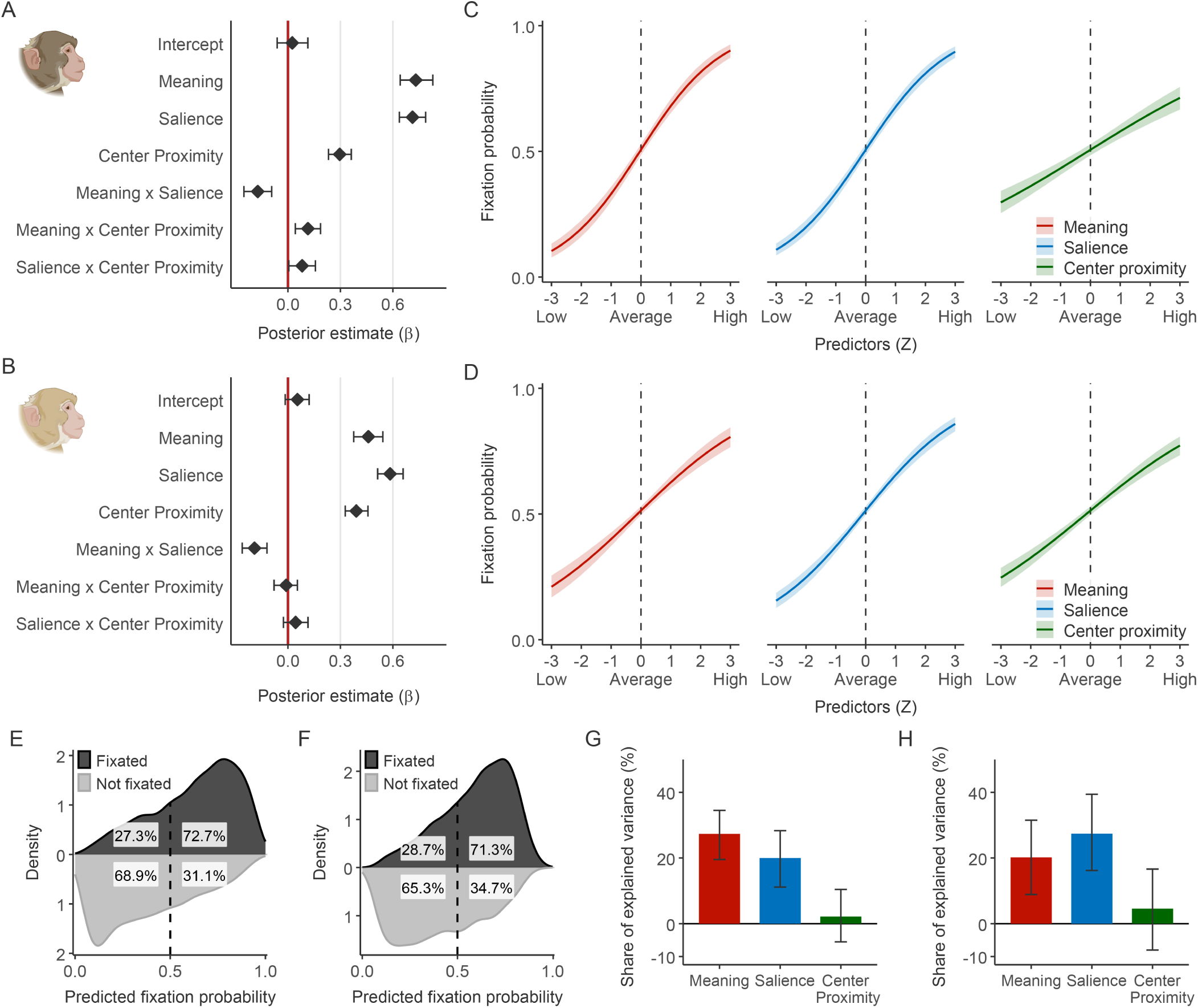
Scene meaning is a robust predictor of fixation selection. (A, B) Posterior estimates of the intercept, main effects, and two-way interactions for Monkey V (A) and Monkey I (B). Points represent the posterior mean of *β* coefficients, and error bars indicate the 95% highest density interval (HDI). The vertical red line marks 0 (no effect). Predictors are considered reliable if their 95% HDI excludes 0. (C, D) Predicted fixation probability as a function of the three main spatial predictors for Monkey V (C) and Monkey I (D). Curves visualize the marginal effects of meaning (red), salience (blue), and center proximity (green) as they vary from *−*3 to +3 standard deviations (*z*-scores), while holding other predictors constant at their means (0). Solid lines indicate the posterior mean prediction, and shaded areas represent the 95% credible interval. (E, F) Model classification performance for Monkey V (E) and Monkey I (F). Density distributions show the model’s predicted probabilities for observed fixations (dark gray) and non-fixated locations (light gray). The vertical dashed line represents the classification threshold (0.5), and the percentages indicate the proportion of observations in each class that fall above or below the threshold. (G, H) Relative importance of spatial predictors for Monkey V (G) and Monkey I (H). Bars represent each predictor’s unique contribution as a percentage of the model’s total explained variance (% Δ*R*^2^) for meaning (red), salience (blue), and center proximity (green). Error bars indicate the 95% highest density interval (HDI). Created in BioRender. Soyuhos, O. (2026) https://BioRender.com/rozb1cy.

**Table 1.**
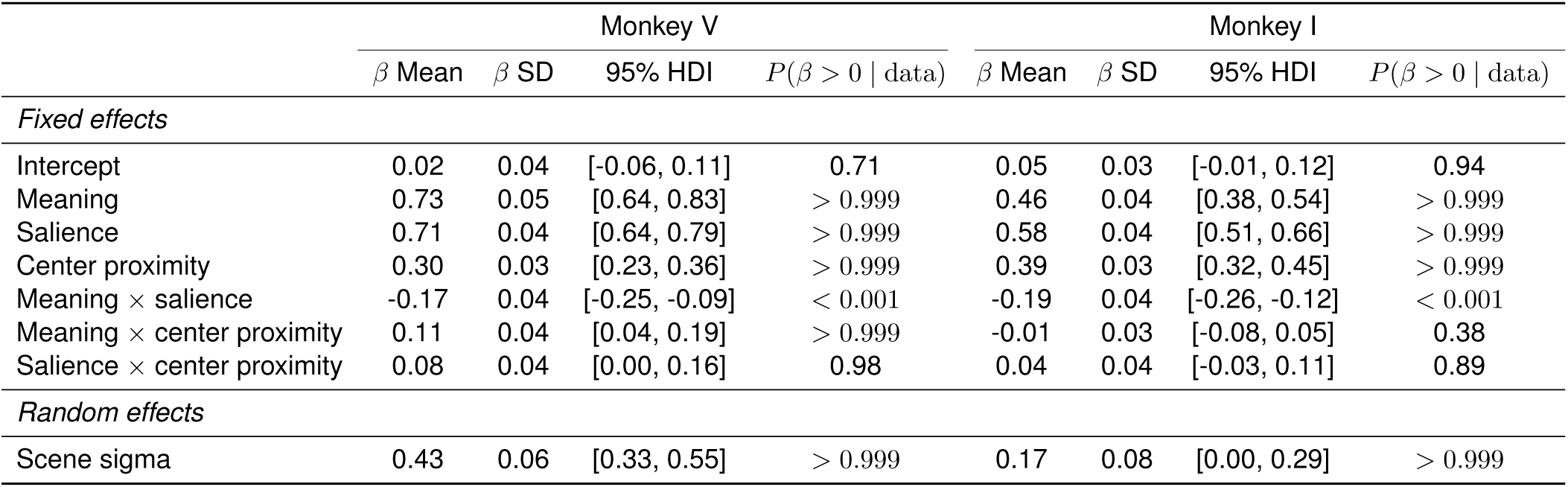
Posterior estimates for the Fixation model. The fixed effects (main effects and two-way interactions) and the scene-level random effect are presented for the model predicting whether a sampled location was fixated. Columns display the posterior mean and standard deviation (SD) of *β* coefficients, the 95% highest density interval (HDI), and the probability of a positive effect (*P* (*β >* 0 *|* data)). Probabilities close to 1 indicate strong evidence for a positive effect, whereas probabilities close to 0 indicate strong evidence for a negative effect. Values above 0.999 or below 0.001 are reported as *>* 0.999 or *<* 0.001 to avoid implying exact certainty. The three-way interaction among meaning, salience, and center proximity was small and did not differ reliably from zero in either animal (Monkey V: *β* = *−*0.02, 95% HDI [*−*0.10, 0.05]; Monkey I: *β* = *−*0.06, 95% HDI [*−*0.13, 0.01]).

|  | Monkey V |  |  |  | Monkey I |  |  |  |
|---|---|---|---|---|---|---|---|---|
|  | β Mean | β SD | 95% HDI | P(β > 0 \| data) | β Mean | β SD | 95% HDI | P(β > 0 \| data) |
| *Fixed effects* |  |  |  |  |  |  |  |  |
| Intercept | 0.02 | 0.04 | [-0.06, 0.11] | 0.71 | 0.05 | 0.03 | [-0.01, 0.12] | 0.94 |
| Meaning | 0.73 | 0.05 | [0.64, 0.83] | > 0.999 | 0.46 | 0.04 | [0.38, 0.54] | > 0.999 |
| Salience | 0.71 | 0.04 | [0.64, 0.79] | > 0.999 | 0.58 | 0.04 | [0.51, 0.66] | > 0.999 |
| Center proximity | 0.30 | 0.03 | [0.23, 0.36] | > 0.999 | 0.39 | 0.03 | [0.32, 0.45] | > 0.999 |
| Meaning × salience | -0.17 | 0.04 | [-0.25, -0.09] | < 0.001 | -0.19 | 0.04 | [-0.26, -0.12] | < 0.001 |
| Meaning × center proximity | 0.11 | 0.04 | [0.04, 0.19] | > 0.999 | -0.01 | 0.03 | [-0.08, 0.05] | 0.38 |
| Salience × center proximity | 0.08 | 0.04 | [0.00, 0.16] | 0.98 | 0.04 | 0.04 | [-0.03, 0.11] | 0.89 |
| *Random effects* |  |  |  |  |  |  |  |  |
| Scene sigma | 0.43 | 0.06 | [0.33, 0.55] | > 0.999 | 0.17 | 0.08 | [0.00, 0.29] | > 0.999 |

We next asked whether meaning guides the first fixations of a trial or whether its influence builds over the viewing period. In humans, meaning-based guidance is present from the earliest fixations (Henderson & Hayes, 2017, 2018). We therefore extended the Fixation model with the order of each fixation in the trial, grouped into fixations 1–3, 4–6, 7–10, and 11 or later (Temporal model). Meaning guided the earliest fixations in both monkeys (fixations 1–3: Monkey V: *β* = 0.73, 95% HDI [0.54, 0.92]; Monkey I: *β* = 0.44, 95% HDI [0.28, 0.60]), and its effect did not change reliably for later fixations (Figure S5). None of the meaning *×* fixation order interactions excluded zero (Monkey V: *β* = *−*0.10 to 0.12; Monkey I: *β* = *−*0.03 to 0.08). Salience also showed no credible change across the trial (Figure S5). Together, these results show that meaning guided fixation selection beyond salience and center bias, from the earliest fixations of a trial.

### Model performance and relative contributions of predictors

We first quantified the overall explanatory power of the predictors (Table S1). For Monkey V, the Fixation model achieved a Bayesian *R*^2^ of 0.26 (95% HDI [0.25, 0.28]) and a point-biserial correlation of *r* = 0.48 (95% HDI [0.479, 0.484]) between the model’s predicted probabilities and the observed fixation behavior. In terms of classification accuracy, the model successfully identified 73% of fixated locations and 69% of non-fixated locations (Figure 2E). For Monkey I, the model achieved a Bayesian *R*^2^ of 0.20 (95% HDI [0.18, 0.21]) and a point-biserial correlation of *r* = 0.43 (95% HDI [0.431, 0.433]), correctly identifying 71% of fixated locations and 65% of non-fixated locations (Figure 2F). We also tested whether the Fixation model extends beyond the scenes and locations it was fitted on. If the relationship between the predictors and gaze is general, a model fitted on some scenes should predict where the animals look in others. In five-fold cross-validation over scenes, each scene was predicted by a model fitted without it. Performance was essentially unchanged: the held-out area under the receiver operating characteristic curve was 0.776 in Monkey V and 0.745 in Monkey I, within 0.003 of the in-sample values (Figure S6). Beyond the sampled locations, the model’s predicted maps correlated with fixation density over the whole image at 0.31 in Monkey V and 0.27 in Monkey I (Figure S7). To put these values in context, we used each animal’s consistency across two viewings of the same scenes as a behavioral reference: the fixation densities of the first and second viewings correlated at 0.36 and 0.42 (Figure S4A). Over the repeated scenes, the model reached 86% of this reference in Monkey V and 65% in Monkey I.

To assess the relative importance of each map, we computed the incremental explained variance (Δ*R*^2^) by refitting reduced models that excluded each predictor and all associated interaction terms, and calculated each predictor’s relative percentage share of the total explained variance (% Δ*R*^2^) (Figure 2G and H). In Monkey V, meaning accounted for a large unique share of variance (Δ*R*^2^ = 0.07, 95% HDI [0.05, 0.09]), representing 27.3% of the model’s total explanatory power (95% HDI [19.5%, 34.5%]). Salience also provided a substantial unique contribution (Δ*R*^2^ = 0.05, 95% HDI [0.03, 0.07]; % Δ*R*^2^ = 20.0% [11.2, 28.3]), while center proximity contributed negligible unique variance (Δ*R*^2^ = 0.006, 95% HDI [*−*0.01, 0.03]; % Δ*R*^2^ = 2.2% [*−*5.5, 10.4]) (Figure 2G). In Monkey I, salience accounted for a large unique share of variance (Δ*R*^2^ = 0.05, 95% HDI [0.03, 0.08]), representing 27.4% of the model’s total explanatory power (95% HDI [16.2%, 39.4%]). Meaning also provided a substantial unique contribution (Δ*R*^2^ = 0.04, 95% HDI [0.02, 0.06]; % Δ*R*^2^ = 20.2% [8.9, 31.5]), while center proximity contributed negligible unique variance (Δ*R*^2^ = 0.01, 95% HDI [*−*0.02, 0.03]; % Δ*R*^2^ = 4.6% [*−*8.0, 16.6]) (Figure 2H). These variance-partitioning analyses therefore indicate that, across both monkeys, meaning-based guidance exerts a robust influence on overt attention after accounting for visual salience and center bias.

The Δ*R*^2^ values above measure only what each predictor explains that the others do not, so variance shared between correlated predictors is attributed to neither. The three predictors are not fully independent. Meaning and salience were correlated in both animals (*r* = 0.44 and 0.42), with weaker correlations between meaning and center proximity (*r* = 0.34 and 0.31) and between salience and center proximity (*r* = 0.19 and 0.18) (Figure S2B and C). We therefore also computed each predictor’s contribution within every model that contains it. Each predictor explained more as the only spatial predictor than its unique contribution in the full model. With center proximity omitted, meaning contributed Δ*R*^2^ = 0.10, 95% HDI [0.08, 0.12] against salience’s 0.05 [0.03, 0.07] in Monkey V and 0.07 [0.05, 0.09] against 0.06 [0.03, 0.08] in Monkey I, a difference of 0.05 [0.03, 0.08] and 0.01 [*−*0.01, 0.04] that excluded zero only in Monkey V (Figure S8). Meaning thus contributed more than salience in Monkey V, while the two contributed comparably in Monkey I.

### Attention prioritizes high-meaning regions, with salience’s role increasing as meaning decreases

We next examined whether meaning and salience combine additively in guiding fixation selection. Under an additive account, salience shifts the log-odds of fixation by the same amount at every level of local meaning. A negative interaction would instead indicate a sub-additive combination, in which salience adds less to fixation probability at locations whose local meaning is already high. We observed a consistent negative interaction between meaning and salience in both monkeys (Monkey V: *β* = *−*0.17, 95% HDI [*−*0.25, *−*0.09], *P* (*β >* 0) *<* 0.001; Monkey I: *β* = *−*0.19, 95% HDI [*−*0.26, *−*0.12], *P* (*β >* 0) *<* 0.001; Table 1). Model-based predictions further quantified this pattern (Figure 3A and B). We calculated the change in predicted fixation probability when image salience shifts from low (*−*2 SD) to high (+2 SD) at different levels of local meaning. At low meaning (*−*2 SD), increasing salience from low to high raised fixation probability by 63 percentage points in Monkey V (95% HDI [53, 73]) and 68 percentage points in Monkey I (95% HDI [60, 77]). In contrast, at high meaning (+2 SD), this change was reduced to 22 percentage points (95% HDI [11, 34]) for Monkey V and 16 percentage points (95% HDI [4, 29]) for Monkey I. Example locations illustrate the same pattern (Figure 3C). In Monkey V, the handles of a pair of scissors (top left) had a predicted fixation probability of 0.57 despite low salience, close to that of high-contrast edges with low meaning (0.55 and 0.56), whereas plain surfaces low in both had 0.12 and 0.14. These results are inconsistent with purely additive integration. Instead, meaning and salience combined sub-additively: high-meaning locations drew fixations even when their salience was low, and salience contributed more to fixation probability as local meaning decreased.

**Figure 3.**
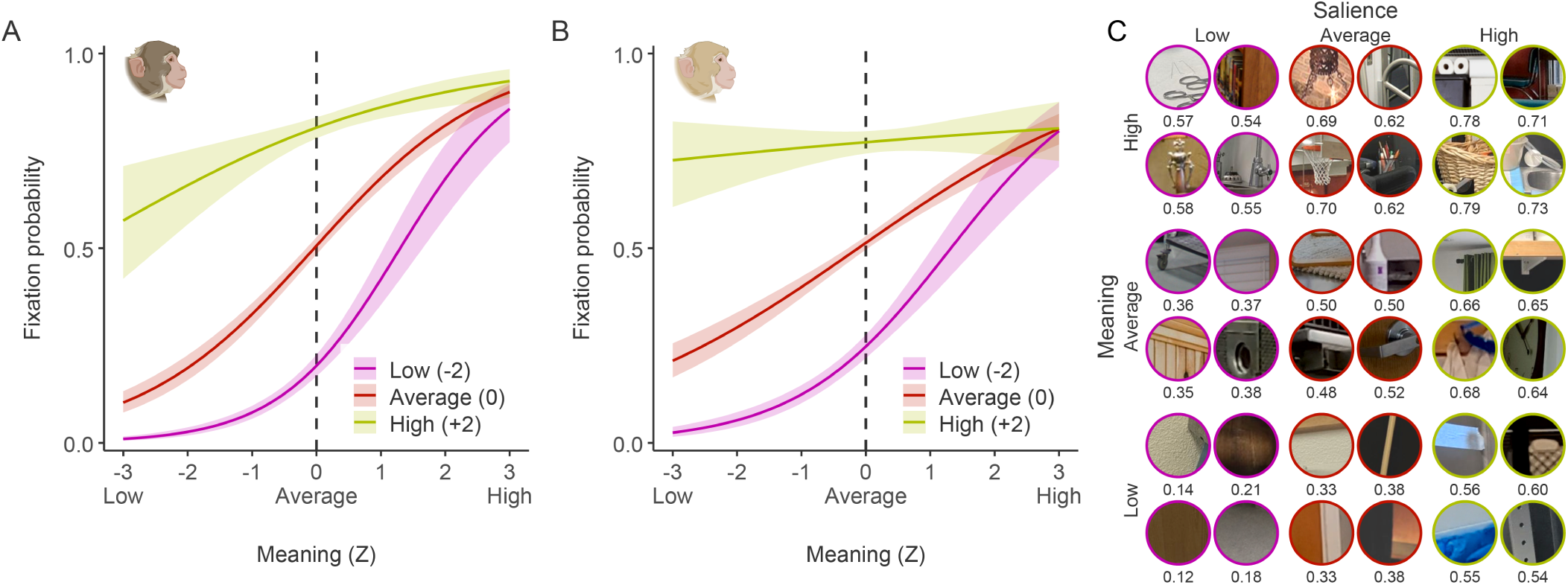
Attention is directed to high-meaning regions, with salience playing an increasing role as meaning decreases. (A, B) Visualization of the meaning *×* salience interaction for Monkey V (A) and Monkey I (B). Curves show the predicted fixation probability as a function of meaning at three representative levels of image salience: low (*−*2 SD, purple), average (0 SD, red), and high (+2 SD, green). Solid lines represent the posterior mean prediction, and shaded areas indicate the 95% credible interval. The vertical dashed line marks the average meaning value (0). In both animals, at locations where local meaning is high, fixation probability remains consistently high with less influence of visual salience. In contrast, at locations where local meaning is low, visual salience strongly modulates fixation probability. (C) Example locations grouped by local meaning (rows) and salience (columns), each divided into the bottom (low), middle (average), and top (high) thirds of each animal’s sampled locations. Each circle shows the 3*^◦^* window over which meaning and salience were averaged, outlined in the salience colors of A and B. In each group, the two examples on the left are from Monkey V and the two on the right from Monkey I. The number below each circle is the predicted fixation probability from that animal’s Fixation model at the window’s meaning and salience, with center proximity at its average. Created in BioRender. Soyuhos, O. (2026) https://BioRender.com/rozb1cy.

### Scene familiarity weakens meaning-based guidance of attention

We further asked whether meaning-based attentional guidance is context-dependent or remains stable across conditions. To test this, we examined whether long-term familiarity with an environment modulates meaning-based guidance. Three competing hypotheses motivated this analysis. First, familiarity might sharpen meaning-based guidance, such that prior knowledge of a scene’s layout drives an even stronger prioritization of high-meaning regions. Second, familiarity might weaken meaning-based guidance, such that knowing the environment reduces the need to exclusively target high-meaning regions, allowing attention to be distributed more frequently to less meaningful areas. Third, the strength of meaning-based attentional guidance might be a fixed trait, operating identically regardless of the observer’s prior experience with the environment.

To distinguish among these hypotheses, we extended the Fixation model to include familiarity (familiar vs. unfamiliar environment; Figure 4A) and its interactions with meaning, salience, and center proximity (Table S2; Familiarity model). Because the familiar and unfamiliar scenes were photographed in different environments, however, the two sets also differ in content. Relative to unfamiliar scenes, familiar scenes were lower in average meaning, spread meaning more evenly within the scene, and were less visually cluttered, while the within-scene correlation between meaning and salience and the within-scene standard deviation of salience did not differ reliably (Figure S9; Table S3). Any effect attributed to familiarity could therefore reflect scene content instead. To separate the two, we fitted the Familiarity model on scenes matched between the two conditions on all five statistics, selected by cardinality matching to a standardized difference below 0.1 (124 of 193 scenes in Monkey V, 118 of 175 in Monkey I), and repeated the analysis at 0.05 as a sensitivity check (Figure S10).

**Figure 4.**
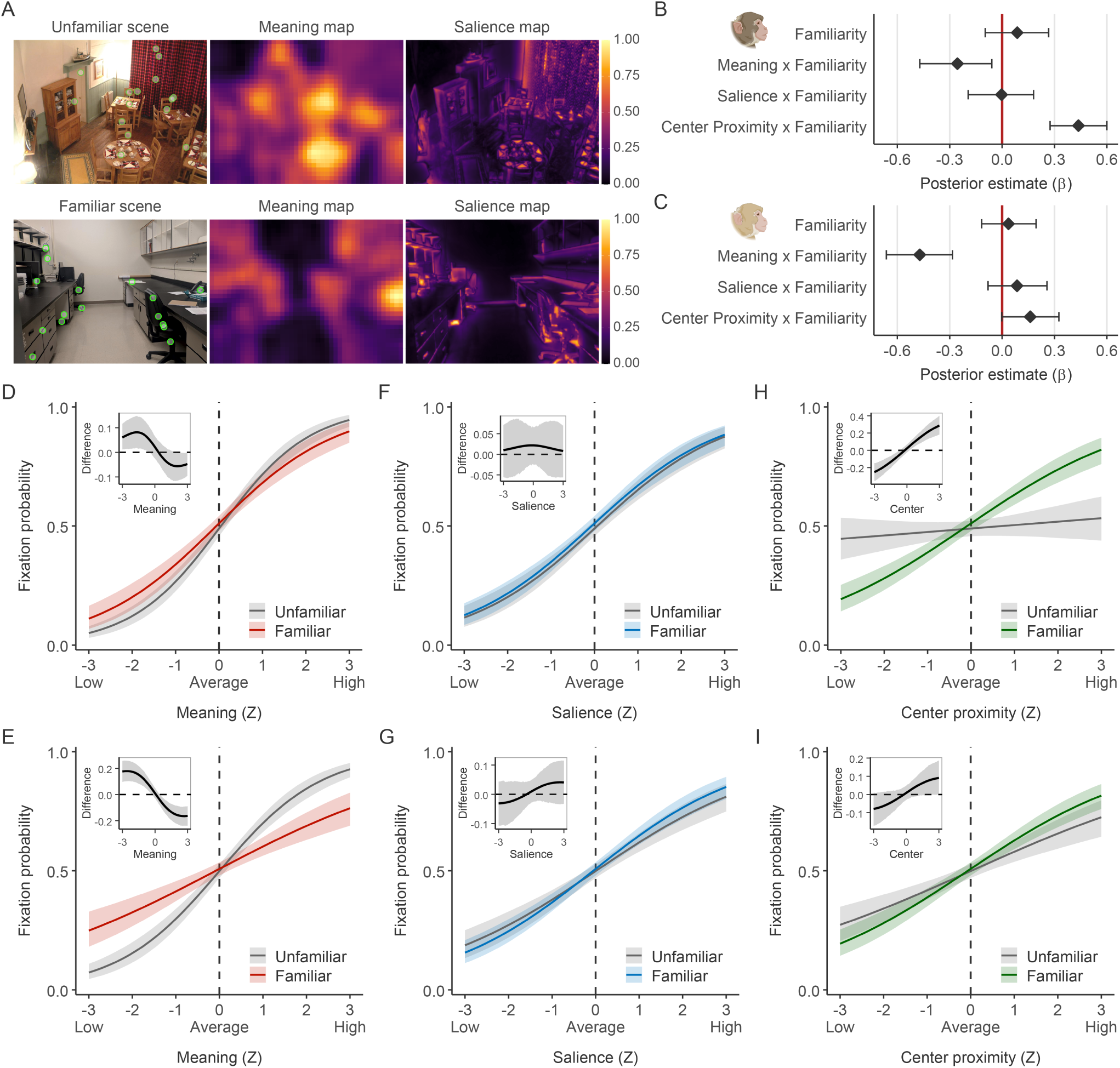
Attentional guidance by meaning is stronger in unfamiliar environments. (A) Example unfamiliar and familiar scenes. The top row displays an unfamiliar scene with fixations (green dots) alongside its meaning and salience maps. The bottom row displays a familiar scene with its corresponding maps. (B, C) Posterior estimates of familiarity effects for Monkey V (B) and Monkey I (C). Points represent the posterior mean of the familiarity coefficient and of its interactions with the spatial predictors; error bars indicate the 95% HDI. The vertical red line marks 0. Note the negative interaction between meaning and familiarity in both animals. (D–I) Predicted fixation probability as a function of spatial predictors for unfamiliar and familiar scenes. Top row (D, F, H) shows results for Monkey V; bottom row (E, G, I) shows results for Monkey I. Curves visualize fixation probability as a function of meaning (D, E), salience (F, G), and center proximity (H, I) for familiar (colored lines) versus unfamiliar (gray lines) scenes. Solid lines represent the posterior mean prediction for each condition, and shaded areas are the 95% credible interval of that prediction. Insets show the posterior mean difference in fixation probability between the two conditions, familiar minus unfamiliar, with its 95% HDI; the dashed line marks 0. In both animals, meaning guided fixation more strongly in unfamiliar than in familiar scenes. All models were fitted on scenes matched between the two conditions on five scene statistics (Figure S9). Created in BioRender. Soyuhos, O. (2026) https://BioRender.com/rozb1cy.

The main effect of familiarity did not differ reliably from zero (Monkey V: *β* = 0.09, 95% HDI [*−*0.10, 0.27]; Monkey I: *β* = 0.04, 95% HDI [*−*0.12, 0.19]) (Figure 4B and C). In both monkeys, however, we observed a reliable negative interaction between meaning and familiarity (Monkey V: *β* = *−*0.26, 95% HDI [*−*0.47, *−*0.06], *P* (*β >* 0) = 0.008; Monkey I: *β* = *−*0.47, 95% HDI [*−*0.66, *−*0.28], *P* (*β >* 0) *<* 0.001), which was retained under the stricter matching criterion (Figure 4B and C; Figure S10). Familiarity reduced the influence of meaning on fixation selection: less meaningful regions received more fixations and more meaningful regions fewer fixations in familiar than in unfamiliar scenes (familiar minus unfamiliar at *−*2 SD of meaning: +0.08, 95% HDI [+0.02, +0.14] in Monkey V and +0.17, 95% HDI [+0.10, +0.25] in Monkey I; at +2 SD: *−*0.06, 95% HDI [*−*0.12, *−*0.002] and *−*0.16, 95% HDI [*−*0.22, *−*0.09]), supporting the hypothesis that familiarity weakens meaning-based guidance (Figure 4D and E). Meaning still guided fixation in familiar scenes in both animals, but less strongly than in unfamiliar scenes (Monkey V: *β* = 0.72, 95% HDI [0.56, 0.87]; Monkey I: *β* = 0.38, 95% HDI [0.26, 0.51]). In contrast, neither monkey showed a reliable interaction between salience and familiarity (Monkey V: *β* = 0.00, 95% HDI [*−*0.19, 0.18]; Monkey I: *β* = 0.09, 95% HDI [*−*0.08, 0.26]) (Figure 4B, C, F, and G). The positive interaction between center proximity and familiarity was strong in Monkey V, with a trend in the same direction in Monkey I (Monkey V: *β* = 0.44, 95% HDI [0.27, 0.60], *P* (*β >* 0) *>* 0.999; Monkey I: *β* = 0.16, 95% HDI [*−*0.002, 0.32], *P* (*β >* 0) = 0.97) (Figure 4B, C, H, and I).

The matched sets were also compared on five low-level image statistics that were not used for matching (luminance, contrast, edge energy, spectral slope, and colorfulness), of which colorfulness differed reliably between conditions in both animals and edge energy in Monkey I alone (Table S3). With colorful-ness added to the model as a covariate, the meaning *×* familiarity interaction was retained in both animals (Monkey V: *β* = *−*0.32, 95% HDI [*−*0.54, *−*0.09]; Monkey I: *β* = *−*0.58, 95% HDI [*−*0.78, *−*0.38]), and the same held with edge energy as the covariate (Monkey V: *β* = *−*0.32, 95% HDI [*−*0.52, *−*0.11]; Monkey I: *β* = *−*0.52, 95% HDI [*−*0.71, *−*0.33]; Table S4). The unique contribution of meaning (Δ*R*^2^) pointed in the same direction, with higher values in unfamiliar than in familiar scenes (Monkey V: 0.09, 95% HDI [0.05, 0.12] versus 0.06 [0.02, 0.09]; Monkey I: 0.08 [0.04, 0.11] versus 0.02 [*−*0.01, 0.06]), whereas that of salience was similar (Monkey V: 0.06 [0.02, 0.09] versus 0.05 [0.01, 0.09]; Monkey I: 0.04 [0.00, 0.07] versus 0.05 [0.01, 0.08]). Together, these results show a common pattern across subjects: familiarity reduced the influence of meaning on fixation selection, whereas the influence of salience did not change reliably.

### Attentional guidance by meaning in relation to scene engagement

Attention not only determines where to look (spatial selection) and what to prioritize (object- or feature-based selection), but also regulates the magnitude of engagement, i.e., the intensity and duration of visual sampling itself. Here, we asked whether scene-by-scene variations in engagement modulate how meaning and salience guide fixation selection. We operationally defined engagement as the total duration of fixations directed at the scene image during each trial. Engagement varied substantially across scenes for both Monkey V (mean = 3586 ms, SD = 547 ms; range = 2015–4463 ms) and Monkey I (mean = 3439 ms, SD = 472 ms; range = 2087–4205 ms) (Figure 5A and B). Importantly, this trial-by-trial variance in engagement was not related to scene familiarity. Total engagement duration did not differ significantly between unfamiliar and familiar scenes for either subject (Wilcoxon rank-sum test: *p* = 0.78 for Monkey V and *p* = 0.52 for Monkey I; Figure S11).

**Figure 5.**
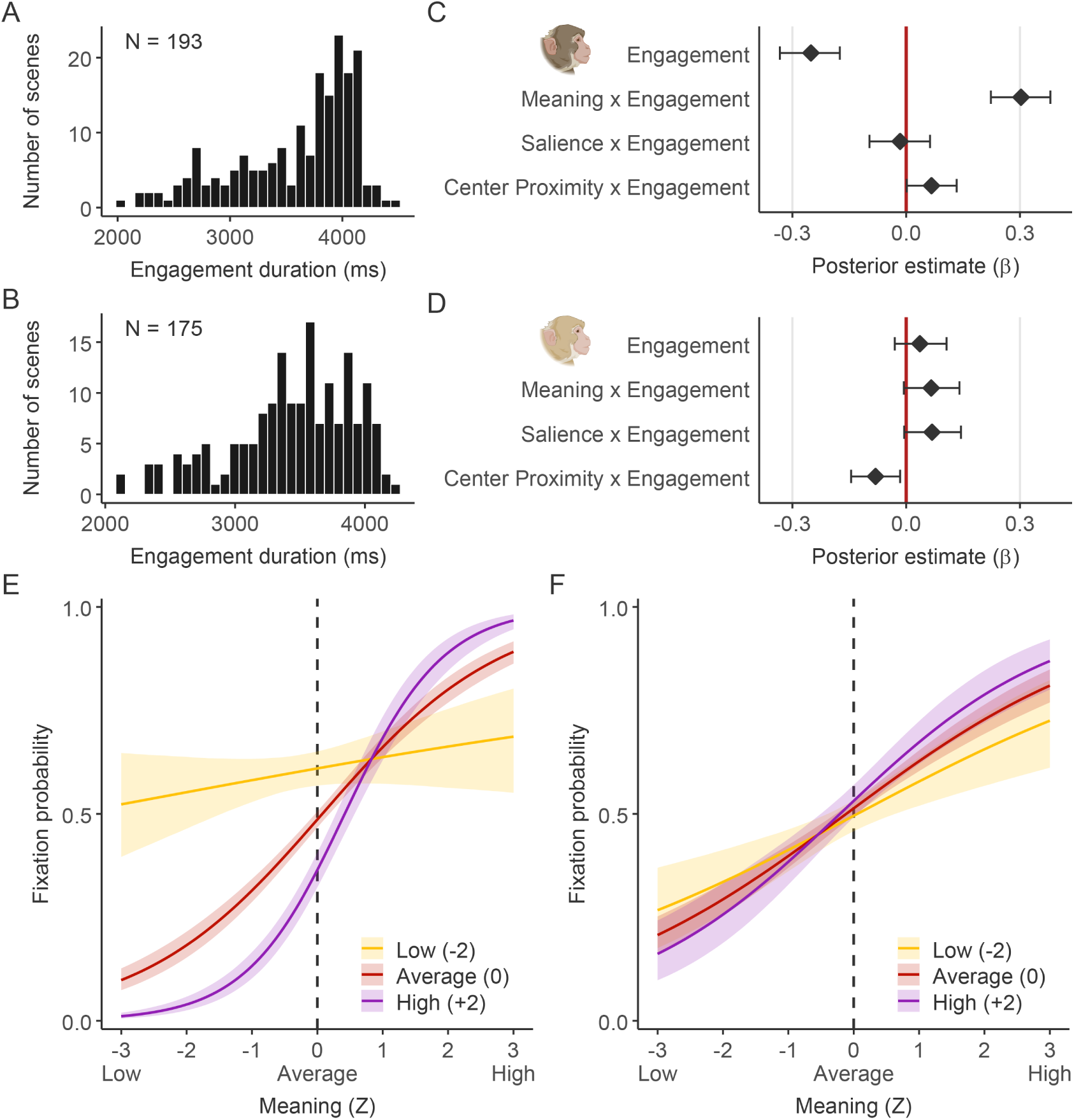
Attentional guidance by meaning in relation to scene engagement. (A, B) Distribution of engagement duration (total fixation time per scene) for Monkey V (A) and Monkey I (B). Histograms show the variability in scene viewing time across trials. (C, D) Posterior estimates of engagement effects for Monkey V (C) and Monkey I (D). Points represent the posterior mean of the engagement coefficient and of its interactions with the spatial predictors; error bars indicate the 95% HDI. The vertical red line marks 0. (E, F) Visualization of the meaning *×* engagement interaction for Monkey V (E) and Monkey I (F). Curves show the predicted fixation probability as a function of meaning at three representative levels of engagement: low (*−*2 SD, yellow), average (0 SD, red), and high (+2 SD, purple). Solid lines represent the posterior mean prediction, and shaded areas indicate the 95% credible interval. Created in BioRender. Soyuhos, O. (2026) https://BioRender.com/rozb1cy.

We extended the Fixation model to include interactions between engagement and the spatial predictors (Table S5; Engagement model). We also fitted this model to both animals together, with monkey as a fixed factor, to estimate the average effect across the two animals and the difference between them. Across the two animals, the effect of meaning on fixation probability increased with higher engagement (*β* = 0.18, 95% HDI [0.13, 0.24], *P* (*β >* 0) *>* 0.999), whereas the effect of salience did not (*β* = 0.03, 95% HDI [*−*0.03, 0.08], *P* (*β >* 0) = 0.83), and the two interactions differed reliably (difference = 0.16, 95% HDI [0.07, 0.24]). This overall pattern came mainly from Monkey V, and the meaning *×* engagement interaction differed between the animals (difference = 0.24, 95% HDI [0.13, 0.34]). For Monkey V, this interaction was substantial (*β* = 0.30, 95% HDI [0.22, 0.38], *P* (*β >* 0) *>* 0.999), indicating that meaning was a stronger determinant of gaze in trials with longer exploration, whereas salience was unaffected (*β* = *−*0.02, 95% HDI [*−*0.09, 0.07], *P* (*β >* 0) = 0.35) (Figure 5C and E). For Monkey I, the interactions with meaning and salience were of the same size, and both intervals included zero (meaning: *β* = 0.07, 95% HDI [*−*0.01, 0.14], *P* (*β >* 0) = 0.96; salience: *β* = 0.07, 95% HDI [*−*0.01, 0.14], *P* (*β >* 0) = 0.96) (Figure 5D and F). These results indicate that, across the two animals, attentional engagement strengthened meaning-based guidance, but that this effect was strong in Monkey V, with a trend in the same direction in Monkey I.

Engagement and fixation count are positively correlated (Monkey V: *r* = 0.79; Monkey I: *r* = 0.64), so the meaning *×* engagement interaction could reflect the number of fixations available rather than engagement itself. We therefore refitted the Engagement model using only the first 10 fixations of each scene, which holds fixation count constant across trials and leaves out late fixations. The interaction persisted across the two animals (*β* = 0.14, 95% HDI [0.08, 0.20]) and in Monkey V (*β* = 0.24, 95% HDI [0.15, 0.33]), whereas its interval included zero in Monkey I (*β* = 0.04, 95% HDI [*−*0.04, 0.13], *P* (*β >* 0) = 0.84) (Figure S12). The number and timing of fixations therefore did not account for the interaction in Monkey V.

Engagement could also reflect the animal’s general state rather than the scene being viewed. We therefore took two measures of the animal’s state before each scene appeared: the time taken to acquire the fixation point and the local rate of trials aborted during the fixation period. These measures explained 4% of the variance in engagement in Monkey V and 3% in Monkey I. We refitted the Engagement model with engagement replaced by its residual from a regression on these measures. The meaning *×* engagement interaction was nearly unchanged (Monkey V: *β* = 0.27, 95% HDI [0.20, 0.35]; Monkey I: *β* = 0.07, 95% HDI [*−*0.01, 0.14], *P* (*β >* 0) = 0.97). Across the two animals, it remained reliable (*β* = 0.17, 95% HDI [0.12, 0.22]) and larger than the salience *×* engagement interaction (difference = 0.14, 95% HDI [0.05, 0.22]). The engagement effect was therefore not explained by these measures of the animals’ general state.

## Discussion

In this study, we asked whether meaning-based guidance of overt attention is uniquely human or extends to non-human primates. We found that scene meaning robustly guided fixation selection in rhesus macaques, with an influence comparable to that of low-level visual salience after accounting for center bias. This guidance was present from the earliest fixations of a trial, and the model generalized to scenes withheld from fitting. These findings provide the first evidence that a human-derived map of scene meaning accounts for where non-human primates look during free-viewing. Moreover, meaning and salience combined sub-additively: high-meaning locations drew fixations even when their salience was low, while visual salience emerged as a stronger guide for eye movements as local meaning decreased. Importantly, meaning-based guidance was not determined solely by image content, but was also modulated by behavioral context. Meaning guided fixation more strongly in unfamiliar than in familiar environments, where less meaningful areas received relatively more fixations. In addition, attentional engagement modulated meaning-based guidance: the influence of meaning strengthened in trials with greater engagement, whereas that of salience showed no consistent change. This effect was strong in Monkey V, with a trend in the same direction in Monkey I. Together, these findings suggest that meaning-based guidance operates alongside visual salience and center bias, thereby expanding current models of primate attention and providing a behavioral foundation for investigating its underlying neural substrates.

### Macaques share the human attentional bias toward informative scene regions

The monkeys preferentially fixated the scene regions that human observers judge informative, indicating that this attentional bias is not unique to humans. This conclusion is consistent with earlier behavioral work. Macaques preferentially orient toward socially relevant stimuli (Deaner et al., 2005; Machado et al., 2011; Solyst & Buffalo, 2014), reward- or aversion-associated objects (Ghazizadeh et al., 2016), and illusory faces in inanimate objects (Taubert et al., 2017), demonstrating that categorical structure and behavioral relevance shape perceptual selection. During search in natural scenes, macaque saccades are also guided by expected reward and by learned associations between targets and scene context (Segraves, 2023; Segraves et al., 2017). In humans, fixations are drawn to objects beyond what feature salience predicts (Einhäuser et al., 2008; Nuthmann & Henderson, 2010), whereas in a direct comparison low-level features accounted for a larger share of macaque than of human fixations (Wilming et al., 2017). Our results extend these findings to general scene viewing, suggesting that the spatial distribution of scene meaning, as judged by human observers, guides macaque attention beyond low-level salience, even in the absence of explicit task demands.

Evidence from human development points the same way. Human infants aged 4 to 12 months preferentially fixate scene regions rated as high in meaning (Oakes et al., 2024). The effect holds after salience and center bias are accounted for, as in the present analysis, and strengthens across the first year of life. Infants lack the knowledge that adult raters bring to the task, so this pattern suggests that their attention operates on object-level representations that develop through visual experience and cortical maturation. Macaques may prioritize the same regions on a similar basis.

The meaning maps used here were produced by DeepMeaning, an image-computable model built on a pretrained vision–language transformer (Hayes & Henderson, 2025). Rather than relying on low-level image statistics, this framework estimates the local meaning of scene regions, such as identifiable objects and their informative relationships. Like the human meaning maps it reproduces, DeepMeaning shows a large drop in estimated meaning when the meaning of a region is removed by diffeomorphic scrambling while its image features are preserved (Hayes & Henderson, 2022a, 2025), so its maps index meaning rather than visual structure alone. Hayes and Henderson (2025) also examined what these estimates contain. They projected the model’s representation of each scene patch onto directions in its vision– language space, each defined by a pair of text prompts describing the presence or absence of a property. Four properties were tested: object density, interaction potential (from manipulable objects such as tools to static surfaces), figure–ground organization, and the consistency of objects with their context. Together they explained about half of the variance in the model’s estimates and in human meaning ratings, and each contributed about a quarter of that explained variance. Although the model was trained on human ratings, its ability to predict macaque gaze behavior suggests that the regions it identifies are prioritized by both species. This is consistent with the similar categorical organization of object representations in human and macaque inferotemporal cortex (Kriegeskorte et al., 2008). Why macaques prioritize these regions is the next question: the regions may be meaningful to the animals in their own terms, for example as objects that afford action, or may draw gaze through the identifiable visual structure that accompanies meaning (see Limitations and future directions).

### Neural substrates for meaning-based attentional guidance

Although our findings are behavioral, they lead to testable predictions for the neural circuits that guide overt visual attention. Dorsal frontoparietal and subcortical structures, including lateral intraparietal cortex (LIP), frontal eye fields (FEF), and superior colliculus, are thought to implement priority maps that determine saccadic targets by integrating both visual salience and behavioral goals (Bisley & Goldberg, 2010; Buschman & Kastner, 2015; Fecteau & Munoz, 2006; Moore & Zirnsak, 2017; Soyuhos et al., 2026b; Xia et al., 2024). Visual salience signals, derived from image-based feature contrasts, are known to contribute to these priority maps (Chen et al., 2020; White et al., 2017). Our results suggest that scene meaning provides an additional source of guidance, which predicts that neurons in these structures carry a signal that covaries with local meaning after visual salience and center proximity are accounted for. The negative interaction between meaning and salience further indicates that the two signals do not combine additively: locations of high local meaning were fixated even when their salience was low, whereas salience guided selection mainly where local meaning was low.

Two complementary accounts could explain how meaning comes to shape where the animal looks. The first concerns the origin of the meaning signal. A plausible source of meaning-related input lies in the ventral visual and scene-selective cortex. In humans, regions such as parahippocampal place area, occipital place area, and retrosplenial cortex encode spatial layout, object ensembles, and contextual structure (Epstein & Baker, 2019; Malcolm et al., 2016; Soyuhos et al., 2026a). Activity in these areas is associated with the meaning of fixated scene regions (Henderson et al., 2020). Macaques have putative homologs of these areas (Nasr et al., 2011). Their inferotemporal cortex contains object- and scene-selective regions (Bao et al., 2020; Bell et al., 2009; Kornblith et al., 2013) and is directly connected with LIP and FEF (Schall et al., 1995; Webster et al., 1994). It is therefore a candidate source of the meaning-related signal predicted above. This pathway devotes specialized areas to objects and scene layout, so informative regions could draw gaze because their content is strongly represented (Xia et al., 2024). The second concerns what the behavior is for. Rather than being drawn passively to over-represented content, the monkeys may be actively sampling their environment for information (Gottlieb, 2012; Gottlieb & Oudeyer, 2018; Gottlieb et al., 2013; Kidd & Hayden, 2015). On this view, meaning-based guidance is one expression of information seeking during natural vision, in which gaze is drawn toward the regions whose representations are most informative to sample (Gottlieb, 2012). The two accounts are not mutually exclusive: a representational bias in the ventral cortex could supply the signal that an information-seeking system evaluates when deciding where to look next. Further experiments will be needed to support or distinguish these possibilities.

### Meaning-based attentional guidance in relation to familiarity and engagement

The influence of meaning on attentional guidance varied with scene familiarity and attentional engagement. Prior work has shown that attention during scene viewing is shaped not only by the physical properties and content of the images, but also by task demands (Borji & Itti, 2014; Castelhano et al., 2009; Yarbus, 1967) and prior experience (Becker & Rasmussen, 2008; Li et al., 2018; Summerfield et al., 2006). Eye movements can reveal memory for previous experience and relational structure, even when explicit report is absent or incomplete (Hannula et al., 2010; Hannula & Ranganath, 2009; Schmidig et al., 2025). In the present study, familiar scenes depicted environments with which the animals had extensive prior experience. Meaning guided fixation more strongly in unfamiliar environments than in these familiar ones, where less meaningful regions received relatively more fixations. This suggests that long-term memory for an environment reduces the influence of meaning, with no reliable change in that of salience. One interpretation is that the monkeys sought information more actively when the environment was novel. In an unfamiliar scene, the content of meaningful regions is still unknown, so fixating them yields the greatest gain in information. In a familiar scene, much of that content is already stored in memory, leaving less to be learned from the same regions and freeing gaze to sample less informative parts of the image. This account is consistent with evidence that novelty itself motivates exploration in primates (Jaegle et al., 2019; Ogasawara et al., 2022) and enhances attentional priority independently of reward (Foley et al., 2014), and with the view that gaze is directed toward locations where uncertainty can be reduced (Gottlieb & Oudeyer, 2018). The selectivity of the effect also fits this account: stored knowledge of a scene should reduce the information to be gained from its meaningful content, but should leave the physical conspicuity of its features unchanged.

Attentional engagement also shaped meaning-based guidance from trial to trial. On trials with higher engagement, fixations tracked the spatial distribution of meaning more closely. A combined model supported this relationship across the two animals. It was reliable in Monkey V, with a trend in the same direction in Monkey I. The effect did not simply follow from engaged trials containing more fixations or unfolding over a longer period: in Monkey V, it was already present within the first 10 fixations of a trial. Engagement itself can arise from several sources, including the animal’s behavioral state and the behavioral relevance of the scene being viewed. Arousal is likely one contributor, but engagement is broader than arousal and may reflect how fully the animal was drawn into viewing a given scene. We could not isolate the arousal component with pupil size, the most common index of arousal, because pupil diameter also varies with luminance and gaze position, both of which changed from scene to scene and throughout free-viewing (Hayes & Petrov, 2016). Instead, we used behavioral measures of the animals’ general state taken before each scene appeared. They explained only a small part of engagement, and the effect was nearly unchanged when this part was removed. More controlled experiments will be needed to separate these contributions directly.

Overall, both animals showed the same pattern of guidance: meaning, salience, and center proximity increased the probability of fixation, meaning and salience combined sub-additively, and familiarity reduced the influence of meaning. The animals differed mainly in magnitude. The meaning coefficient was smaller in Monkey I than in Monkey V (*β* = 0.46 versus 0.73), as was the modulation by engagement. These differences could reflect prior experience, as Monkey V had a considerably longer history of laboratory training, but with two subjects the contribution of experience cannot be separated from that of individual traits such as temperament. Human observers likewise differ reliably in how strongly faces, text, and other categories attract their gaze (de Haas et al., 2019).

### Limitations and future directions

Several limitations bear on the interpretation of these results. First, the evidence is correlational. We modeled the association between map values and fixation selection in intact scenes and did not manipulate the scenes, so the finding that meaning predicts fixation selection beyond salience and center proximity is a model-based association rather than a demonstration that meaning guides attention causally. In humans, this question has been addressed with manipulations that alter scene meaning while preserving local image features, such as scene inversion (Hayes & Henderson, 2022b) and diffeomorphic scrambling of local content (Hayes & Henderson, 2022a), and applying these manipulations to macaque viewing is a direct next step. Second, the meaning maps are derived from human judgments. Our results show that macaques, like humans, prioritize the regions that human observers judge informative, but not that they do so for the same reasons. A macaque is unlikely to know what a telephone is for, but it may treat one as an object to grasp or explore, which would make such regions meaningful to the animal in its own terms. The same regions also contain identifiable objects and visual structure that could attract gaze on their own. The manipulations above would help separate these accounts, as would maps built by rating scene patches for what they afford, such as graspability (Rehrig et al., 2020), and testing whether relatedness in meaning between objects predicts gaze transitions between them, as reported for human scene inspection (Hwang et al., 2011). Recording human observers viewing the same scenes would also allow the two species to be compared directly, including whether there are regions of high meaning that humans fixate but macaques do not.

In addition, fixation location is the endpoint of both attentional prioritization and saccade programming, so our analysis does not separate the two, and the priority-map account offered above is an interpretation of the behavioral pattern rather than a conclusion drawn from it. Moreover, the familiar scenes were limited to the vivarium and laboratory spaces available to photograph, so they also differed in content from the unfamiliar scenes, a difference we reduced by matching the two sets on five content statistics. Finally, our salience control covers only low-level feature contrast. Deep saliency models also learn mid- and high-level features (Hayes & Henderson, 2021), and how these features relate to meaning and to macaque fixation selection is a question for future work.

## Materials and methods

### Subjects

The experiments involved two healthy male rhesus macaques (*Macaca mulatta*): Monkey V and Monkey I. At the time of data collection, Monkey V was 14 years old and weighed 13.3 kg, and Monkey I was 14 years old and weighed 13.8 kg. Both animals were sourced from the California National Primate Research Center and housed in standard primate facilities with environmental enrichment, in accordance with institutional guidelines. The monkeys were maintained on a controlled fluid access schedule to ensure motivation for the juice reward. Daily health and hydration status were monitored. All procedures were approved by the University of California, Davis Institutional Animal Care and Use Committee (Protocol #24082) and were conducted in accordance with the NIH Guide for the Care and Use of Laboratory Animals.

### Experimental setup and eye tracking

Animals were seated in a primate chair with the head stabilized using a surgically implanted head-post. The chair was positioned in a sound-attenuated, dimly lit booth facing a computer monitor. Visual stimuli were presented on a 32” LCD monitor (Display++, Cambridge Research Systems) with a resolution of 1920 *×* 1080 pixels and a refresh rate of 120 Hz. The physical width of the display was 70 cm, and the viewing distance was 80 cm from the subject’s eyes. The screen background was a uniform mid-gray (RGB: 128, 128, 128). Stimulus presentation and experimental control were implemented in MATLAB R2023a (MathWorks) using Psychtoolbox-3. Juice reward was delivered through a tube positioned at the subject’s mouth and controlled by a syringe pump (NE-4000-US, New Era Pump Systems Inc.).

Eye position was recorded from the right eye using a video-based infrared eye tracker (EyeLink 1000 Plus, SR Research) sampling at 1000 Hz. The tracker was configured to record gaze position, pupil diameter, and event markers for fixations, saccades, and blinks using the manufacturer’s standard settings. At the beginning of each session, a horizontal–vertical 9-point calibration was performed on a mid-gray background using the standard EyeLink calibration routine, and the display coordinates in the EyeLink system were matched to the Psychtoolbox display window. Raw EyeLink data files (.edf) were imported into MATLAB R2023a using the Edf2Mat toolbox (developed by Etter and Biedermann, University of Zurich).

### Stimuli, design, and behavioral task

Stimuli consisted of 200 naturalistic indoor scenes (1440 *×* 1080 pixels) presented in a randomized (shuffled) design. We did not include humans or other animals in any image to keep the scenes emotionally neutral. The stimulus set included 100 unfamiliar scenes and 100 familiar scenes. Unfamiliar scenes depicted environments the monkeys had not directly experienced prior to the experiment, such as restaurants, bedrooms, kitchens, and offices. Familiar scenes were photographs taken in the monkeys’ own living and experimental environment, including vivarium housing and laboratory spaces, and therefore reflected locations with which the animals had long-term experience. Familiarity was defined by this exposure history and no separate recognition test was administered. Trials were interleaved such that unfamiliar and familiar scenes appeared in a random order, so that familiarity was not confounded with position in the session. Each scene was presented twice, once in each of two blocks separated by a break of about 15 minutes. The order was shuffled once for each animal and the same order was used in both blocks, so every scene occupied the same serial position in both. The two animals therefore saw the scenes in different orders. Each block began with 10 practice trials, which were not analyzed. Unless stated otherwise, all analyses use the first presentation of each scene.

Each trial began with a central black fixation dot (0.5*^◦^* radius) on a mid-gray background. The fixation window was a 6*^◦^ ×* 6*^◦^* square centered on the dot (*±*3*^◦^* horizontally and vertically). In the first phase, the subject had up to 5000 ms to acquire fixation; once acquired, it had to be maintained continuously for 500 ms. If the subject failed to hold fixation, the trial was aborted and repeated after a 1000 ms inter-trial interval (ITI). In the second phase, the fixation dot disappeared and a naturalistic scene was presented at the center of the screen for 5000 ms. During this viewing period, the subject was free to move its eyes and explore the scene. In the third phase, a juice reward was delivered immediately following the viewing period, followed by a 1000 ms ITI.

### Meaning, salience, and center proximity maps

For each scene, we generated three spatial predictor maps at the native image resolution of 1440 *×* 1080 pixels. A meaning map quantified the spatial distribution of local informativeness across each scene, estimated with DeepMeaning (Hayes & Henderson, 2025), an image-computable model of human meaning maps (Henderson & Hayes, 2017). Human meaning maps are produced by dividing a scene into a dense array of overlapping circular patches at two spatial scales (3 and 7 degrees of visual angle), asking naive raters to judge how informative or recognizable each patch is on a six-point scale when it is presented in isolation and in random order, and averaging the ratings at each location. DeepMeaning reproduces these maps from the image alone. Square patches of the scene are passed through a Contrastive Captioner (CoCa) vision–language transformer pretrained on more than two billion image–text pairs, whose weights are frozen, and a linear readout of its features is trained to predict the human meaning map value of each patch. The readout was trained on human meaning maps of 282 scenes from 100 categories with leave-one-scene-out cross-validation, separately for indoor and outdoor scenes. On held-out indoor scenes, the predicted maps correlated with the human maps at *r* = 0.87, the level at which independent groups of human raters agree with each other, and they predicted human fixations as well as the human maps do. The model is never trained on fixation data. We used the indoor model to produce a map for each scene; map values are on the scale of the human ratings and are comparable across scenes. Applying these maps to macaque viewing does not assume that the animals share human semantic knowledge. It asks whether the spatial distribution of regions that human observers judge informative or recognizable predicts where macaques look.

A salience map was computed for each scene using the classic Itti–Koch model (Itti et al., 1998) as implemented in the pySaliencyMap package (github.com/akisatok/pySaliencyMap), which combines multiscale feature maps for luminance contrast, color, and orientation into a single salience map that is normalized within each scene. We chose this model rather than graph-based alternatives to minimize center bias that would otherwise be highly collinear with our center proximity predictor.

A center proximity map represented spatial bias toward the image center and was operationalized as the normalized inverse Euclidean distance from the image center. For a pixel located at coordinates (*x, y*), it was defined as:

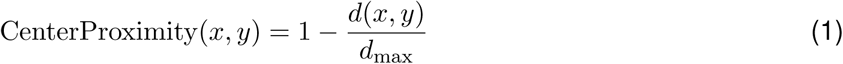

where *d*(*x, y*) is the Euclidean distance from the image center and *d*_max_ is the maximum distance from the center to any corner of the image.

### Fixation data, trial exclusion, and non-fixated samples

For each trial, fixation events (onset time, duration, and *x*–*y* coordinates) were extracted from the Eye-Link data files. The first fixation that began after scene onset was excluded because its location was constrained by the central fixation at trial onset. Fixations were included only if they began within the 5 s free-viewing window and fell within the boundaries of the displayed image. Because the 1440 *×* 1080 image was centered on a 1920 *×* 1080 display, this excluded fixations directed at the gray background to either side. It removed 1.9 fixations per trial for Monkey V and 2.3 fixations per trial for Monkey I, corresponding to 10% and 12% of the fixations recorded during the viewing window.

Prior to model fitting, we applied exclusion criteria to ensure that only trials reflecting active exploration were included. A trial (scene) was excluded from analysis if the subject made fewer than 10 fixations or if the total fixation duration was less than 2000 ms. Both criteria counted only fixations on the image, after the first fixation was excluded (Figure S1). One interrupted trial in Monkey I’s first viewing was also excluded. This resulted in the exclusion of 7 scenes for Monkey V and 24 of the remaining 199 scenes for Monkey I, leaving a final dataset of 193 and 175 scenes, respectively. Most excluded scenes failed both criteria (5 of 7 in Monkey V and 14 of 24 in Monkey I); the remainder had fewer than 10 fixations (2 and 7) or less than 2000 ms of fixation (0 and 3). Excluded trials contained about half as many fixations on the image as retained trials, with fixations of similar duration, indicating that the animal spent less of the viewing window looking at the image. Retained trials were comparable between animals in fixation count (means of 16.3 and 16.8) and total fixation time (3586 *±* 547 and 3439 *±* 472 ms). The final dataset included a total of 3140 fixations for Monkey V and 2932 fixations for Monkey I.

We modeled fixation probability with balanced datasets consisting of fixated locations and matched non-fixated locations (Hayes & Henderson, 2022b; Oakes et al., 2024). For each real fixation (Fixated = 1), we generated one corresponding non-fixated sample (Fixated = 0). To ensure that non-fixated samples represented locations the animal did not fixate, they were drawn from random coordinates within the image boundaries, constrained to be at least approximately 3 degrees of visual angle away from all fixation locations in that scene (Figure 1B). The value of each predictor at a sampled location was taken as the mean of its map within a window of approximately 3 degrees of visual angle centered on that location. The exclusion distance matches the diameter of this window, so that the sampling windows of fixated and non-fixated locations do not overlap. This 3-degree scale also corresponds to the resolution at which meaning is measured, since human meaning maps are rated on circular scene patches at 3 and 7 degrees of visual angle (Henderson & Hayes, 2017) and DeepMeaning is trained to reproduce them. To assess how this choice affects the estimates, we refitted the Fixation model with the scale reduced to approximately 2 degrees and to approximately 1 degree, applied to the window and the exclusion distance alike. The fixated locations and all other analysis steps were unchanged (Figure S3).

### Bayesian generalized linear mixed model

All models were fitted using brms (version 2.22.0; Bürkner, 2017) with a Bernoulli likelihood and a logit link function. We used the default priors provided by brms, which consist of flat priors on regression coefficients and weakly informative priors on the intercept and the scene-level standard deviation. Each model was estimated using four Markov chain Monte Carlo (MCMC) chains with 6000 draws per chain (2000 warm-up, 4000 sampling), resulting in 16000 posterior samples. All models included a random intercept for Scene to capture baseline heterogeneity in fixation likelihood across scenes. Posterior predictions were obtained by evaluating the fitted models on grids of predictor values and averaging over posterior samples. Because each fixation was paired with one non-fixated sample, these predicted probabilities are conditional on the balanced sampling scheme: they give the probability that a location with given feature values belongs to the fixated rather than the non-fixated set, and are reported as relative, model-based effect sizes rather than as absolute probabilities of fixating an arbitrary scene location. Posterior distributions of model parameters, and of quantities derived from them such as Δ*R*^2^ and differences in predicted probability, are summarized by their mean and 95% highest density interval (HDI). The bands around predicted probability curves are 95% equal-tailed credible intervals. Unlike the HDI, an equal-tailed interval is unchanged by the logit transform, so the band on the probability scale is the transformed interval of the underlying linear predictor.

We examined the dependence among the GLMM predictors. Variance inflation factors were low for both subjects (maximum VIF = 1.33), but the predictors are not fully independent: meaning and salience correlate at *r* = 0.44 and 0.42 and meaning and center proximity at *r* = 0.34 and 0.31, while salience and center proximity are largely unrelated (*r* = 0.19 and 0.18) (Figure S2). We also inspected the correlations between the posterior parameter estimates, which index how well the model separates the effects of the predictors rather than how independent the predictors themselves are. These were modest for both Monkey V (maximum *|r|* = 0.39) and Monkey I (maximum *|r|* = 0.27; Table S6). Additionally, all fixed effects demonstrated stable convergence (*R*^^^ = 1.00) and high effective sample sizes (ESS *>* 5000) in the Fixation, Temporal, Familiarity, and Engagement models (Table S7).

### Model specifications

For each monkey, continuous predictors (meaning, salience, and center proximity) were *z*-scored across the sampled locations, that is, the fixated locations and their matched non-fixated locations, so that each predictor had a mean of 0 and a standard deviation of 1 before model fitting. Coefficients are therefore expressed per standard deviation of each predictor over the sampled locations. Using these standardized predictors, we addressed our research questions with three main models. First, we fitted a Fixation model to test the effects of meaning, salience, and center proximity, and of their interactions on overt attention:

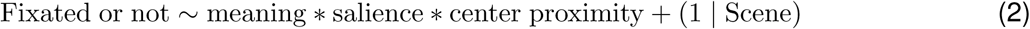

With the predictors standardized over the whole image instead, the Fixation model gave nearly the same coefficients (*β* = 0.74 and 0.49 for meaning and 0.70 and 0.58 for salience).

To examine how guidance changed over the course of a trial, we also extended the Fixation model with the order of each fixation in the trial (Temporal model):

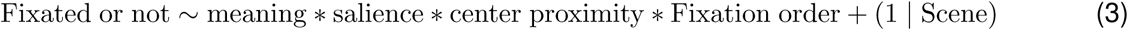

Fixation order grouped the retained fixations by their ordinal number in the trial into four levels: 1–3, 4–6, 7–10, and 11 or later. Fixation 1 was the first fixation retained after the exclusions described above, and fixations 1–3 formed the reference level. Each non-fixated sample took the order of the fixation it was paired with, so every level kept one non-fixated sample per fixation. A single model was fitted across levels, so that the scene intercept was estimated from all samples of a scene, as in the Fixation model.

Second, we fitted a Familiarity model by adding a categorical predictor for familiarity to examine how long-term environmental familiarity alters attentional guidance. Familiarity encoded whether the scene was familiar (Familiarity = 1) or unfamiliar (Familiarity = 0) and was included in the interaction terms:

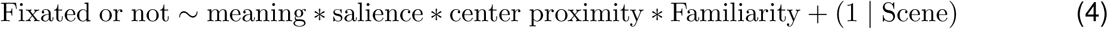

Because the familiar and unfamiliar scenes differ in content as well as in familiarity, we fitted the Familiarity model on a subset of scenes matched between the two conditions on five statistics: the mean and within-scene standard deviation of meaning, the within-scene standard deviation of salience, visual clutter (Feature Congestion; Rosenholtz et al., 2007), and the within-scene correlation between meaning and salience (Fisher *z* transformed). Each was computed over the entire scene. Scene-level average meaning could be matched because meaning ratings are absolute and comparable across scenes, unlike the salience maps, which are normalized within each scene.

Matched sets were selected by cardinality matching (Zubizarreta et al., 2014), implemented in MatchIt 4.7.2 (Ho et al., 2011) with the HiGHS solver (Huangfu & Hall, 2018). The method finds the largest subsample in which the two conditions satisfy specified balance constraints. We required familiar and unfamiliar scenes to be equal in number and the mean of each condition to lie within 0.1 standardized units of the mean of all analyzed scenes on all five statistics simultaneously, and repeated the analysis at 0.05 as a sensitivity check. This retained 124 of 193 scenes for Monkey V and 118 of 175 for Monkey I at the 0.1 criterion, and 118 and 116 at 0.05. The largest residual difference between conditions was 0.08 at the 0.1 criterion and 0.04 at 0.05, below the 0.1 conventionally taken as negligible imbalance (Austin, 2011). The matched scenes were taken from the standardized data described above, so that coefficients remain on the same scale as those of the model fitted on all scenes.

Five further image statistics were computed for each scene but not matched on: mean relative luminance (sRGB values linearized and combined with Rec. 709 weights), RMS contrast (the standard deviation of luminance divided by its mean), edge energy (the mean gradient magnitude of the luminance plane), spectral slope (the rate at which amplitude falls with spatial frequency on log–log axes, averaged over orientations), and colorfulness (the spread and mean of pixel values along the red–green and yellow– blue opponent axes; Hasler & Süsstrunk, 2003). Familiar and unfamiliar scenes were compared on all ten statistics, in all analyzed scenes and in the matched sets at the 0.1 criterion, by standardized difference and Wilcoxon rank-sum test (Table S3). Colorfulness, the only statistic that still differed reliably between conditions in both animals after matching, and edge energy, which still differed in Monkey I, were each added to the primary matched Familiarity model as a covariate in a separate fit. Each covariate was *z*-scored across the sampled locations in the matched scenes and entered as a main effect and in two-way interactions with the three spatial predictors (Table S4):

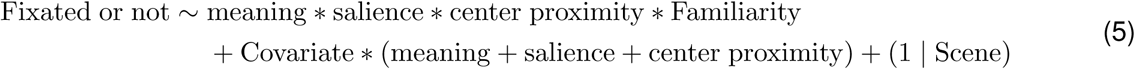

Third, we fitted an Engagement model to investigate how guidance varies with engagement. Engagement was defined as the total duration of fixations directed at the scene image during the 5 s free-viewing period, after the exclusions described above, capturing trial-to-trial fluctuations in the intensity of visual exploration. Fixations accounted for 87% (Monkey V) and 81% (Monkey I) of the viewing window and saccades for a further 13% and 18%, with blinks contributing 0.2% and 3%. Blinks are recorded within saccade events and are therefore included in the saccade percentages. Time classified as neither fixation nor saccade averaged 38 ms and 40 ms per trial. Engagement was compared between unfamiliar and familiar scenes with a Wilcoxon rank-sum test (Figure S11), then *z*-scored and included as an interaction term in the Engagement model:

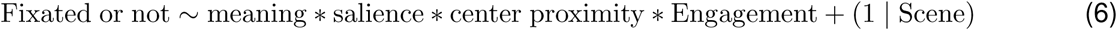

To verify that the engagement interactions did not arise from differences in the number of fixations available per trial, we refitted the Engagement model using only the first 10 fixations of each scene. This subset retained the matched non-fixated sample belonging to each fixation. We also fitted the Engagement model to both animals together, with monkey as a fixed factor coded +0.5 (Monkey V) and *−*0.5 (Monkey I) and crossed with all terms, and with a separate scene intercept for each animal. The main terms of this model give the average of the two animals, and the interactions with monkey give the difference between them. Monkey was not treated as a random effect, because two animals are too few to estimate the variance between them. The same model was fitted to the first 10 fixations of each scene.

We also tested whether the engagement interactions reflected the animals’ general state rather than engagement with the scene. For each trial, we computed two measures of state before scene onset. Fixation acquisition time was the interval between the onset of the fixation point and the entry of gaze into the fixation window. The local abort rate was the proportion of trials in which fixation was acquired and then broken during the 500 ms fixation period. It was computed over the 50 surrounding trials in which fixation was acquired, 25 before and 25 after, leaving out the trial itself. Across the analyzed trials, the median fixation acquisition time was 138 ms in Monkey V and 118 ms in Monkey I, and the local abort rate ranged from 2% to 22% and from 15% to 48%. For each animal, engagement was regressed on log-transformed fixation acquisition time and the local abort rate. The Engagement model and the model of both animals were then refitted with engagement replaced by the residuals of this regression, *z*-scored in the same way. The residuals were nearly identical for windows of 20 to 50 trials (*r ≥* 0.996).

### Model performance and variance partitioning

We quantified the explanatory power of the fixed effects in each model with the Bayesian *R*^2^, calculated using the bayes_R2 function in brms (Gelman et al., 2019). We reported the Bayesian *R*^2^ for the fixed effects, which estimates the proportion of variance in the observed outcome explained by the model’s expected predictions, excluding random scene intercepts. We also calculated the point-biserial correlation between the model’s predicted probabilities and the observed binary outcome. This correlation was computed for each posterior draw and summarized using the mean and 95% HDI. Classification accuracy was the proportion of fixated and of non-fixated locations identified correctly, with a location assigned to the fixated set when its predicted probability exceeded 0.5.

Five-fold cross-validation over scenes tested how well the Fixation model generalizes beyond the scenes it was fitted on. Scenes were assigned at random to five folds of approximately equal size. For each fold, the model was refitted on the remaining scenes with the same specification and used to predict the outcome at the sampled locations of the held-out scenes, so that every location was predicted once by a model that had not been fitted on its scene. Because each scene was presented once per block, a held-out scene is also a held-out trial. Predictions used the fixed effects only, and the in-sample values were computed the same way. Performance was summarized by the area under the receiver operating characteristic curve (Figure S6).

As a behavioral reference for model performance, we quantified within-subject fixation reliability across the two viewings of each scene. To ensure comparability with the GLMM analysis, we applied the same trial-exclusion criteria to the second block and restricted the repeatability analysis to scenes that passed them in both blocks, resulting in 159 repeated scenes for Monkey V and 148 repeated scenes for Monkey I. For each repeated scene, we generated smoothed fixation density maps for the initial and repeated viewings by binning fixation locations into 40 *×* 40 pixel bins (approximately 1 degree) and applying Gaussian smoothing with a full width at half maximum of 3 degrees of visual angle. Bins near the image border receive only part of the smoothing kernel, so each map was divided by the kernel weight that fell inside the image; without this correction the border of every map is attenuated in the same way in both viewings, which inflates the correlation between them. We then calculated the Pearson correlation between the two density maps for each scene. The average correlation across repeated scenes provided a model-free estimate of within-subject repeatability.

To compare the model against this reference on the same scale, we also evaluated the Fixation model over the whole image. The posterior mean fixed effects were applied to the three predictors at every location of each scene, averaged over the same 3-degree window used for the sampled locations and binned to the resolution of the density maps, and the resulting map of predicted fixation probability was correlated with the observed fixation density map of the first viewing of that scene. The scene random intercept was omitted, since it is constant within a scene and does not affect the spatial pattern. For the comparison with the reference, the model’s correlations were averaged over the repeated scenes only.

In a model-free analysis over the whole image, the meaning and salience maps of each scene were evaluated on the grid of the density maps, as the mean map value in the 3-degree window averaged within each bin, and correlated with the observed fixation density map of the first viewing (Henderson & Hayes, 2017). We also computed the semipartial correlation of fixation density with each map, using the residual of that map after a linear regression on the other map and the center proximity map. Means across scenes are reported with 95% confidence intervals from 10000 bootstrap resamples of scenes, each distribution was tested against zero with a one-sample Wilcoxon signed-rank test, and meaning and salience were compared with a paired Wilcoxon signed-rank test.

We calculated the change in Bayesian *R*^2^ (Δ*R*^2^) by comparing the full Fixation model to reduced models in which a specific predictor and all its associated interactions were removed. The distribution of Δ*R*^2^ values was computed from the posterior samples, providing a direct estimate of the incremental explanatory contribution of meaning, salience, and center proximity, respectively. We also calculated each predictor’s unique contribution as a percentage of the full model’s explanatory power (% Δ*R*^2^), summarized in the same way. Because Δ*R*^2^ attributes variance shared between correlated predictors to neither, we also computed each predictor’s contribution to the model’s Bayesian *R*^2^ within every subset of the spatial predictors that contains it, as in dominance analysis (Azen & Budescu, 2003; Figure S8). In addition to the full Fixation model, this required a model for each of the six smaller subsets of the three spatial predictors, each retaining the interactions among the predictors it contains.

To ask whether meaning and salience contributed differently in the two familiarity conditions, we computed a familiarity-split Δ*R*^2^. A single Familiarity model was fitted on the scenes matched at the 0.1 criterion, together with two reduced versions omitting meaning and salience respectively, each with all associated interactions removed. Bayesian *R*^2^ was then evaluated separately on the familiar and the unfamiliar rows of the data, and Δ*R*^2^ taken as the difference between the full and reduced model within each condition.

## Data and code availability

All data and code needed to reproduce the results are publicly available. The Zenodo deposit (https://doi.org/10.5281/zenodo.24639928) contains the scene images and their meaning and salience maps, the per-scene image statistics, the raw eye-tracking recordings, the fixation and model input tables, the per-trial summaries, and the fitted models. The deposit excludes the images and maps of the 100 familiar scenes, which show the monkeys’ vivarium and laboratory spaces. These files are available from the corresponding author on request. The code at https://github.com/orhansoyuhos/ Meaning-based-guidance-of-attention-in-rhesus-monkeys generates fixation density maps and model predictions for any scene and includes instructions for reproducing every figure, table, and reported statistic.

Salience maps were computed with the pySaliencyMap implementation of the Itti–Koch model (github.com/akisatok/pySaliencyMap). Meaning maps were computed with the publicly released Deep-Meaning model (Hayes & Henderson, 2025). We provide an open-source application for computing meaning maps (github.com/orhansoyuhos/deep-meaning-app).

## Acknowledgments

This work was supported by the National Science Foundation under grant 2152260 (NSF NRT Neu-ralStorm) (to O.S., W.H., and B.S.), National Institutes of Health grants EY014924 (to X.C.) and R01EY027792 (to J.M.H.), and the Brain and Behavior Research Foundation (to X.C.).

## Contributions

Study conceptualization: X.C., O.S., J.M.H., and T.R.H.; Experimental design: O.S. and X.C.; Data collection: O.S., T.P.H., B.S., and W.H.; Methodology: O.S., X.C., and T.R.H.; Exploratory analysis: O.S., T.R.H., and W.H.; Formal analysis: O.S.; Visualization: O.S.; Writing – original draft: O.S. and X.C.; Writing – review & editing: O.S., X.C., J.M.H., and T.R.H.; Funding acquisition: X.C., O.S., W.H., and J.M.H.

## Corresponding author

Correspondence to Xiaomo Chen

## Ethics declarations

The authors declare no competing interests.

