## Supplementary information for "Meaning-based guidance of attention in rhesus monkeys during naturalistic scene viewing"

#### Figures

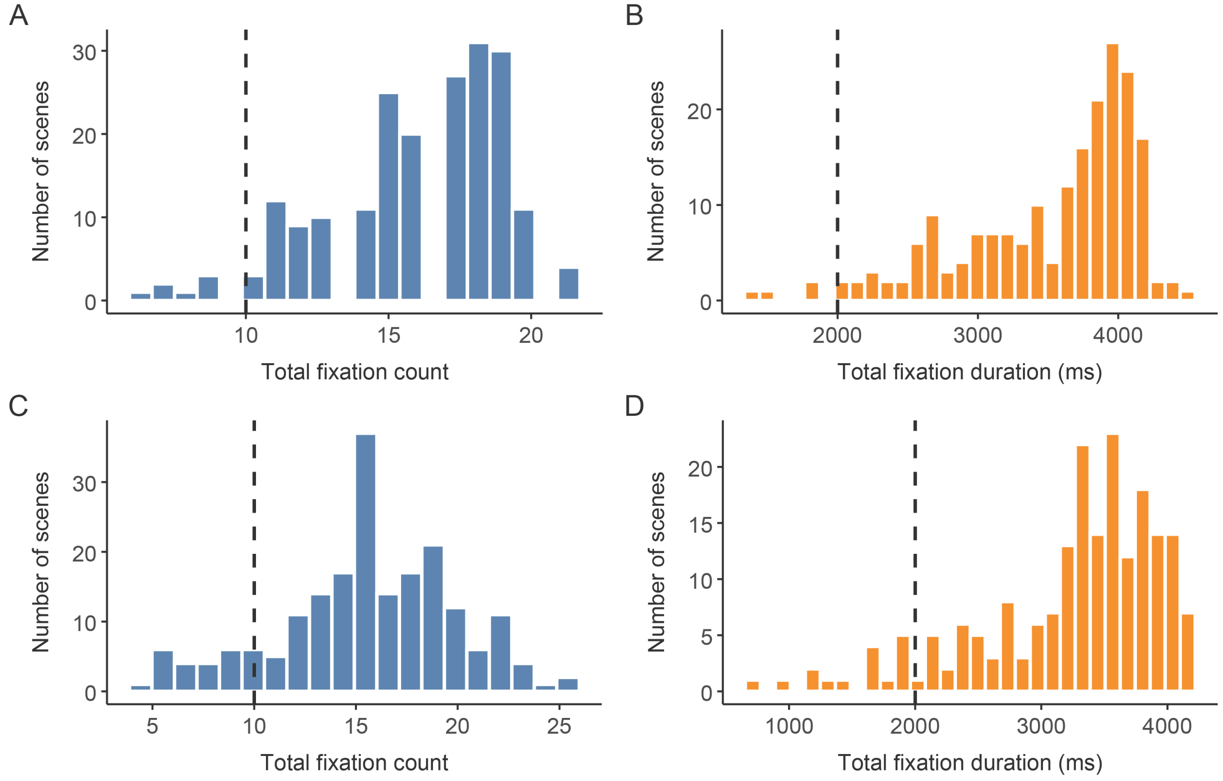


**Figure S1 | Data exclusion criteria.**

(A, B) Histograms showing the distribution of total fixation counts (A) and total fixation duration (B) across all 200 scenes of the first viewing block for Monkey V. (C, D) Corresponding distributions for Monkey I. Vertical dashed lines indicate the thresholds applied after removal of the first fixation after scene onset: scenes were excluded if the subject made fewer than 10 fixations (left column) or if the total accumulated fixation duration was less than 2000 ms (right column). Scenes falling below either threshold were excluded: 7 of 200 for Monkey V and 24 of 199 for Monkey I.

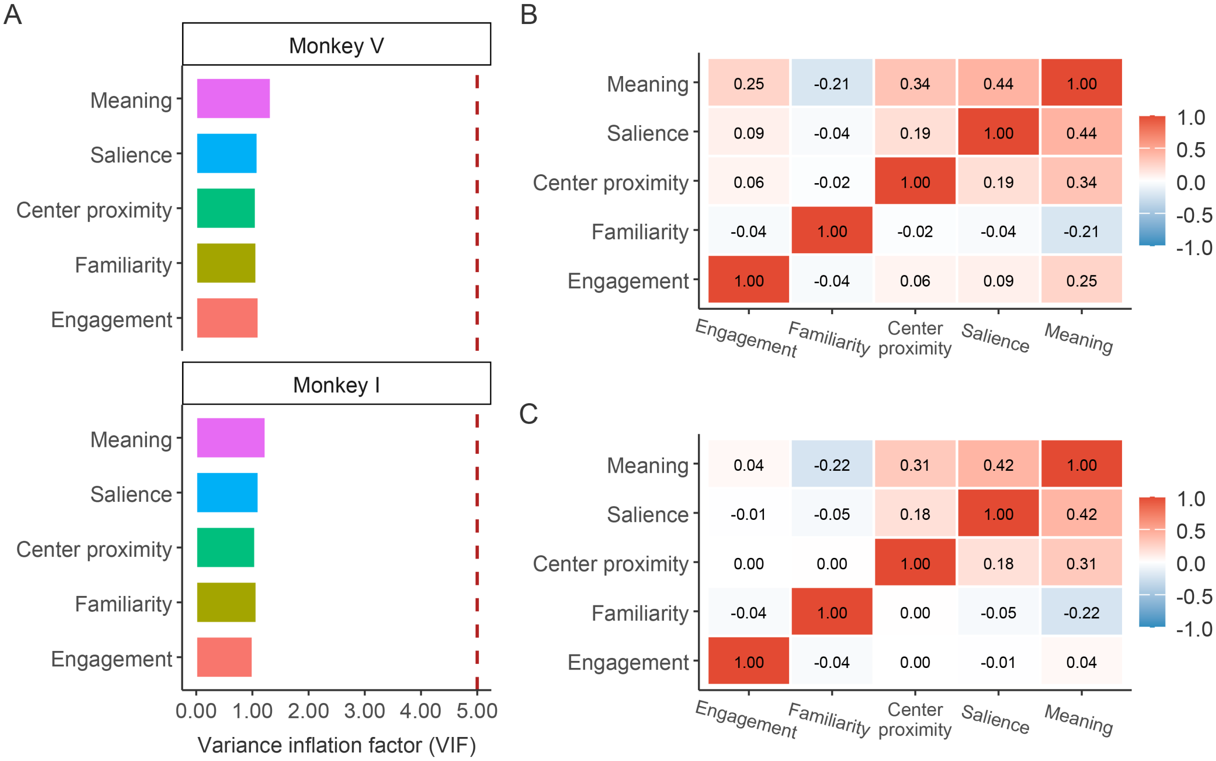


**Figure S2 | Multicollinearity diagnostics and pairwise correlations for model predictors.**

Diagnostics were computed for the fixated and matched non-fixated locations used to fit the generalized linear mixed models. (A) Variance inflation factor (VIF) for the main fixed-effect predictors, calculated separately for Monkey V (top) and Monkey I (bottom). The red dashed line denotes a standard VIF threshold of 5. All predictors exhibit VIF values near 1, indicating that the dependence among them does not inflate the coefficient estimates. (B, C) Pairwise Pearson correlation matrices for the predictors in Monkey V (B) and Monkey I (C): meaning, salience, center proximity, familiarity, and engagement. Cell values and colors represent the correlation coefficient ($r$) between each pair of variables. Among the three spatial predictors, meaning and salience are the most strongly correlated pair in both subjects ($r = 0.44$ and 0.42), followed by meaning and center proximity ($r = 0.34$ and 0.31) and salience and center proximity ($r = 0.19$ and 0.18). The predictors are therefore related but not redundant, and the variance shared between meaning and salience is partitioned in Figure S8.

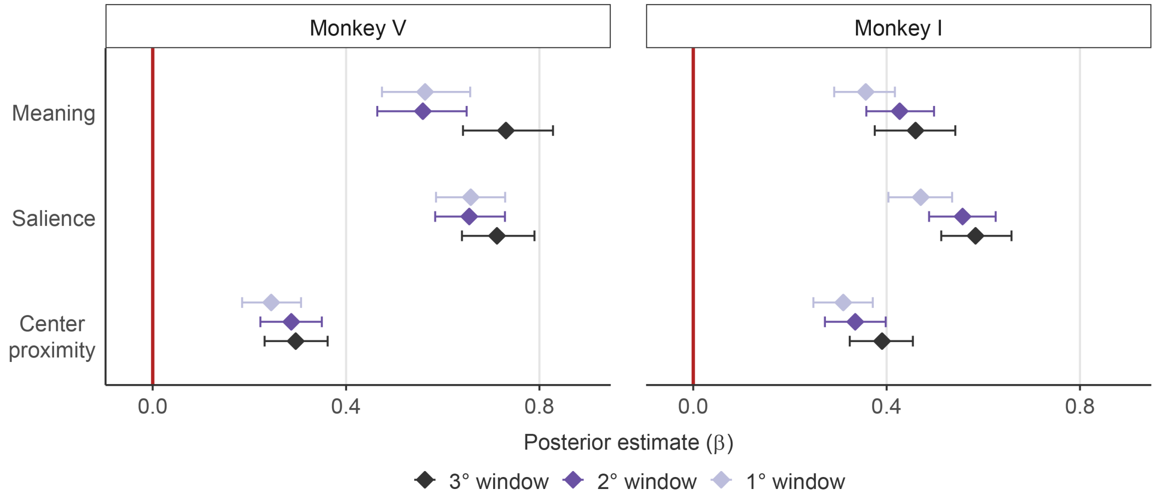


**Figure S3 | Sensitivity of the Fixation model to the spatial scale of sampling.**

In the main analysis, predictor values were averaged over an approximately 3 degree window and each non-fixated location was drawn at least 3 degrees from every fixation in the scene. The sensitivity analyses reduced this scale to 2 and to 1 degree. The exclusion distance equals the window diameter at every scale, so the sampling windows of fixated and non-fixated locations never overlap. Posterior estimates of the three spatial predictors, for Monkey V (left) and Monkey I (right). Diamonds represent the posterior mean and horizontal bars the 95% HDI, in black for 3 degrees and in progressively lighter purple for 2 and 1 degrees. The vertical red line marks 0.

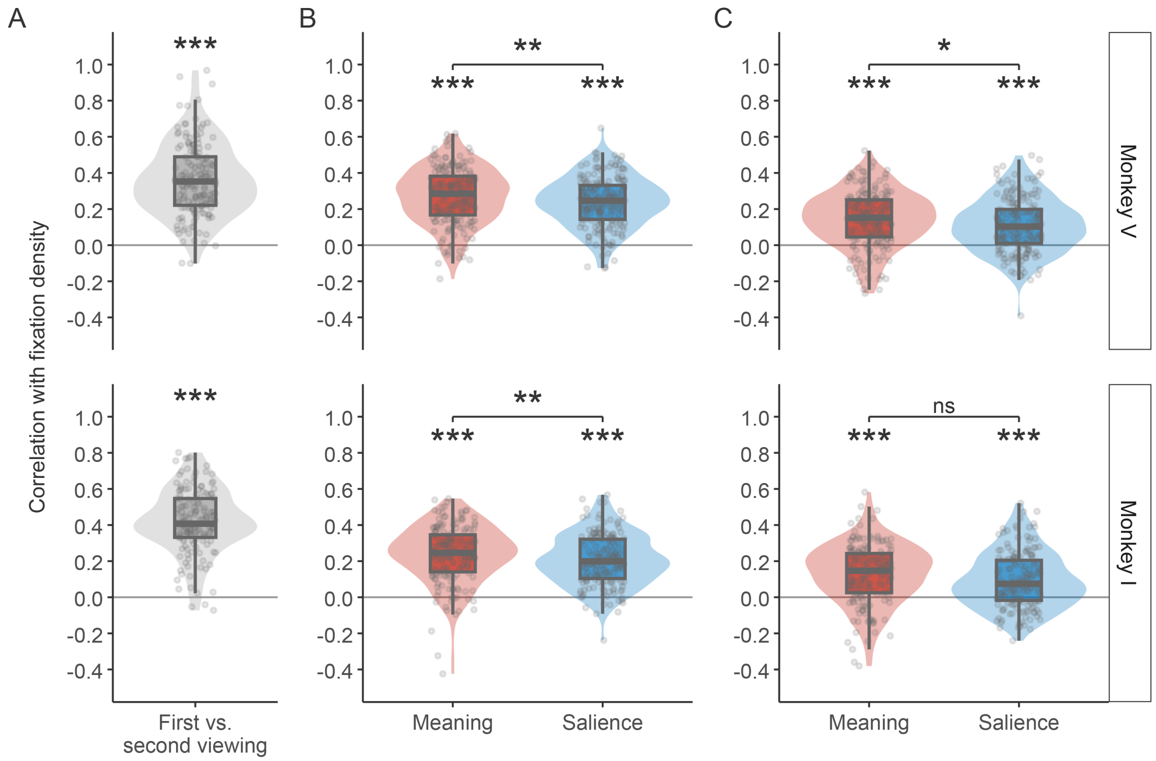


**Figure S4 | Fixation density over the whole image.**

Each point is one scene, and each value is a correlation with the fixation density map of the first viewing of that scene, computed without sampled locations or a model, for Monkey V (top) and Monkey I (bottom). (A) Correlation between the fixation density maps of the first and second viewings of the same scene (159 and 148 scenes). (B) Correlation with the meaning map (red) and with the salience map (blue) (193 and 175 scenes). (C) The same correlations after the other map and center proximity were regressed out of each map (semipartial correlation). Violins show the distribution across scenes, and boxes show the median and interquartile range, with whiskers extending to 1.5 times the interquartile range. Asterisks directly above a distribution mark a one-sample Wilcoxon signed-rank test against zero, and asterisks above a bracket mark a paired Wilcoxon signed-rank test of meaning against salience across scenes (*, $p < 0.05$; **, $p < 0.01$; ***, $p < 0.001$; ns, not significant). Means with 95% confidence intervals for Monkey V and Monkey I: (A) 0.36 [0.33, 0.39] and 0.42 [0.39, 0.45]; (B) meaning 0.27 [0.25, 0.29] and 0.24 [0.21, 0.26], salience 0.24 [0.22, 0.26] and 0.21 [0.19, 0.23]; (C) meaning 0.14 [0.12, 0.17] and 0.13 [0.10, 0.15], salience 0.11 [0.09, 0.13] and 0.09 [0.07, 0.12].

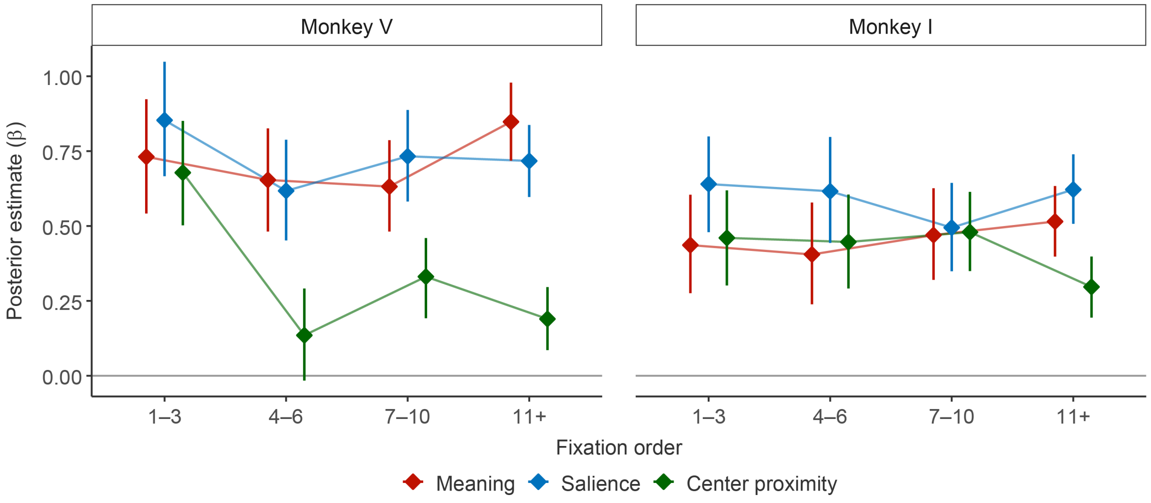


**Figure S5 | Attentional guidance across the course of a trial.**

Posterior estimates of meaning, salience, and center proximity for each group of fixations in the Temporal model, which extends the Fixation model with the order of each fixation in the trial (fixations 1–3, 4–6, 7–10, and 11 or later), for Monkey V (left) and Monkey I (right). Fixations are numbered after the exclusion of the first fixation after scene onset. Diamonds represent the posterior mean and vertical bars the 95% HDI, with meaning in red, salience in blue, and center proximity in green. The horizontal gray line marks 0.

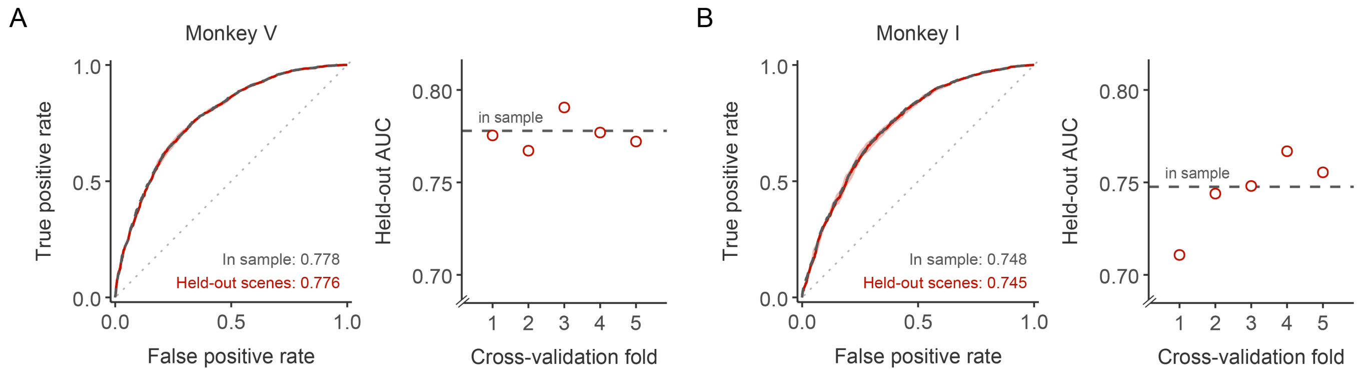


**Figure S6 | Prediction of fixation selection on data the model has not seen.**

(A) Monkey V. Left, receiver operating characteristic (ROC) curves showing how well the model tells fixated locations from their matched non-fixated controls: the fraction of fixated locations correctly identified against the fraction of non-fixated locations wrongly identified. A model with no information follows the dotted diagonal, and the further a curve bows toward the upper left, the better it separates the two classes. The dashed gray curve is the model fitted to all analyzed scenes of the first viewing block and evaluated on those same locations, the fit reported in the main text. The solid red curve is its cross-validated counterpart: the scenes were split into five folds, the model was refitted on four folds at a time, and every location was predicted once by a model that had never seen its scene. The red band is the standard error across the five folds. AUC, the area under the curve, is the probability that a fixated location receives a higher predicted value than a non-fixated location; 0.5 is chance and 1 is perfect separation. Right, the held-out AUC of each fold (red) against the in-sample AUC over all scenes (dashed line). (B) The same for Monkey I.

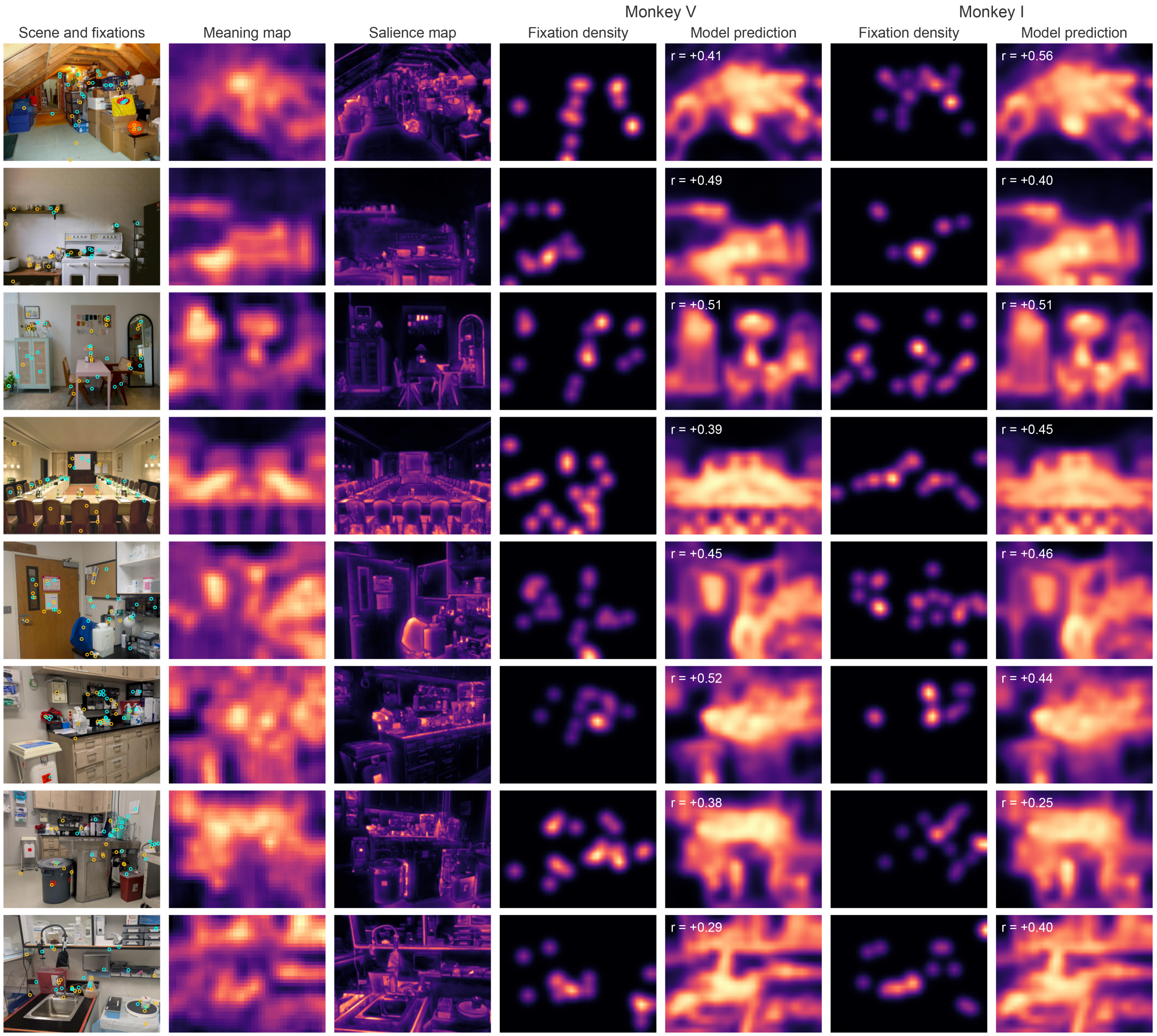


**Figure S7 | Example scenes: fixation density and the model's prediction.**

Eight scenes are shown one per row: four unfamiliar (rows 1 to 4) and four familiar (rows 5 to 8). From left to right: the scene with the fixations of both animals overlaid (Monkey V in yellow, Monkey I in cyan), its meaning map, its salience map, and for each animal the observed fixation density beside the map of fixation probability predicted by the Fixation model. The value inset in each predicted map is the Pearson correlation between that predicted map and that animal's fixation density for that scene. Across all analyzed scenes, this correlation averaged 0.31 in Monkey V and 0.27 in Monkey I. Predicted maps were obtained by applying the posterior mean fixed effects at every location, with each predictor averaged over the same approximately 3-degree window used for the sampled locations. Meaning and salience maps are shown at native resolution; the model averages each within the 3-degree window. All fixations are from the first viewing of each scene.

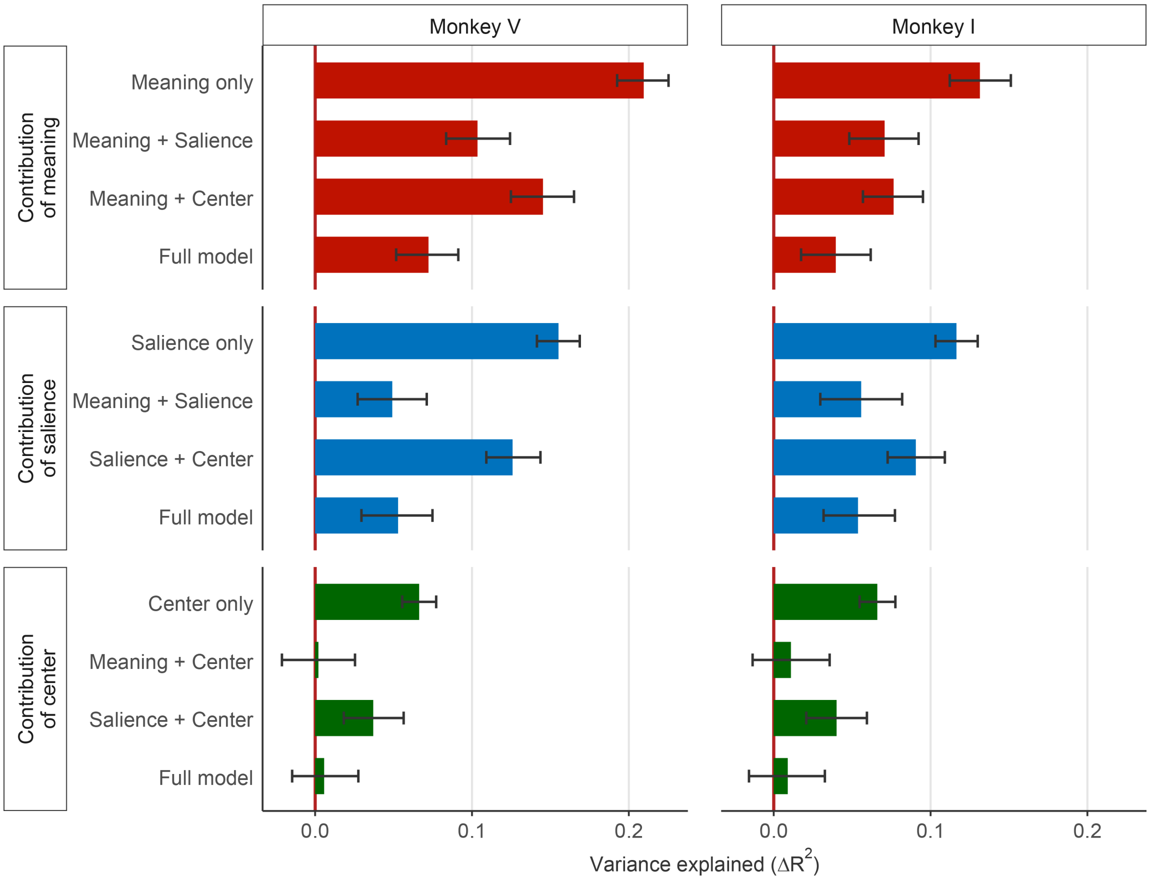


**Figure S8 | Contribution of each spatial predictor across model subsets.**

Each bar gives the difference between a model's Bayesian $R^2$ and that of the same model with one predictor removed, computed for every subset of the three spatial predictors. Columns are the two animals, and rows correspond to the predictor whose contribution is shown, as labeled on the left. Within a row, bars are labeled by the model they refer to, ranging from the model containing that predictor alone to the full model with all three. In the first, that predictor is the only one present, so the bar is that model's whole $R^2$, the proportion of variance in fixation selection it explains. In the last, the other two predictors remain, so the bar is the unique contribution that Figure 2G and H reports as percentages. The contributions of meaning, salience, and center proximity are shown in red, blue, and green respectively. Bars represent the posterior mean, horizontal lines the 95% HDI, and the vertical red line marks 0.

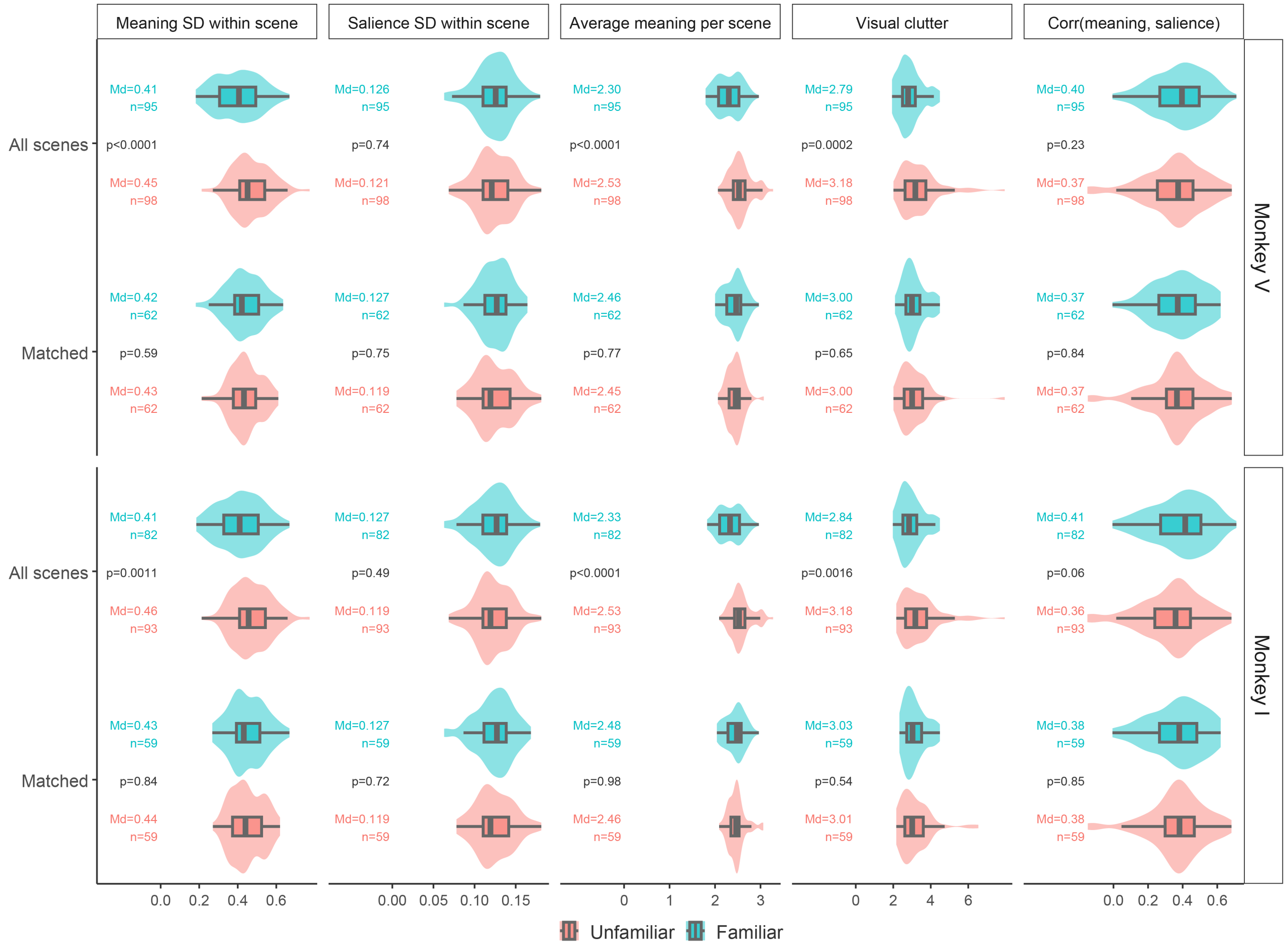


**Figure S9 | Matching the familiar and unfamiliar scene sets on five content statistics.**

Distributions of each statistic in familiar (cyan) and unfamiliar (red) scenes, for Monkey V (top) and Monkey I (bottom). Within each animal, the upper pair of distributions shows all analyzed scenes and the lower pair the subset retained after matching. Each statistic is computed over the entire scene. Columns, left to right: the within-scene standard deviation of meaning, the within-scene standard deviation of salience, the average meaning of the scene, visual clutter based on feature congestion, and the within-scene Pearson correlation between meaning and salience. Because meaning ratings are on an absolute scale that is comparable across scenes, the average meaning of a scene itself can be matched alongside the within-scene statistics, unlike the average salience of a scene. Violins show the kernel density. Boxes show the interquartile range and median. Sample size ($n$) and median (Md) for each group, together with the Wilcoxon rank-sum test between them ($p$), are shown in each panel. Matched sets were built by cardinality matching, which selects the largest subsample in which the mean of each condition lies within a specified standardized distance of the mean of all analyzed scenes on all five statistics simultaneously. Here a threshold of 0.1 retained 124 of 193 scenes in Monkey V and 118 of 175 in Monkey I. This is the set on which the models in Figure 4 were fitted.

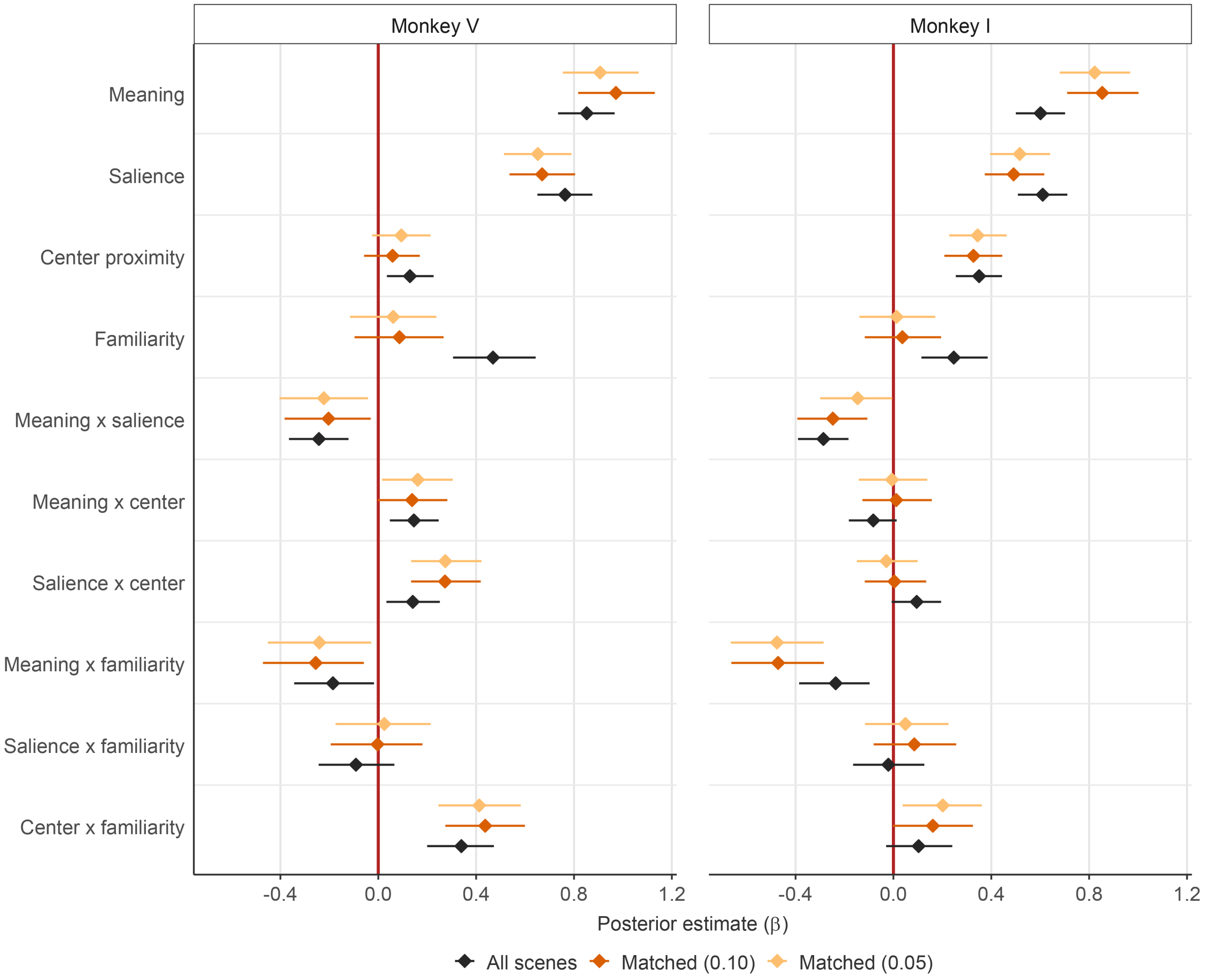


**Figure S10 | Fixed effects of the Familiarity model before and after content matching.**

Each row is one fixed effect of the model (main effects and two-way interactions) and each panel one monkey. Diamonds mark the posterior mean and horizontal lines the 95% HDI. The vertical red line marks zero. Matched sets were selected by cardinality matching, which finds the largest subsample in which the mean of each condition lies within a given standardized distance of the mean of all analyzed scenes on all five scene statistics simultaneously: the average meaning of the scene, the within-scene standard deviation of meaning and of salience, visual clutter, and the within-scene correlation between meaning and salience. Three fits are shown per animal: all analyzed scenes (black; 193 scenes in Monkey V, 175 in Monkey I), scenes matched to a standardized difference below 0.1 (dark orange; 124 and 118 scenes; reported in Figure 4), and scenes matched under a stricter threshold of 0.05 (light orange; 118 and 116 scenes). The model specification, sampling settings, and predictor standardization are identical across the three fits.

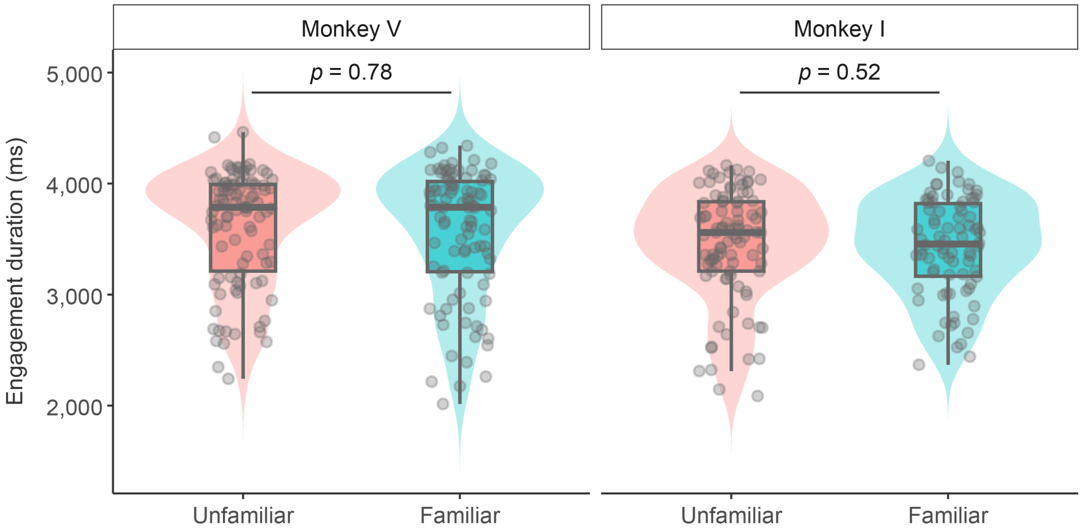


**Figure S11 | Attentional engagement duration does not differ with scene familiarity.**

Violin plots illustrate the distribution of scene-level engagement durations (ms) across unfamiliar (red) and familiar (cyan) scenes for Monkey V (left) and Monkey I (right). Each scatter point represents the total engagement duration for a single trial. Overlaid box plots denote the median (central horizontal line), interquartile range (box edges), and 1.5× interquartile range (whiskers). Engagement duration did not differ significantly between unfamiliar and familiar scenes in either subject (Wilcoxon rank-sum test).

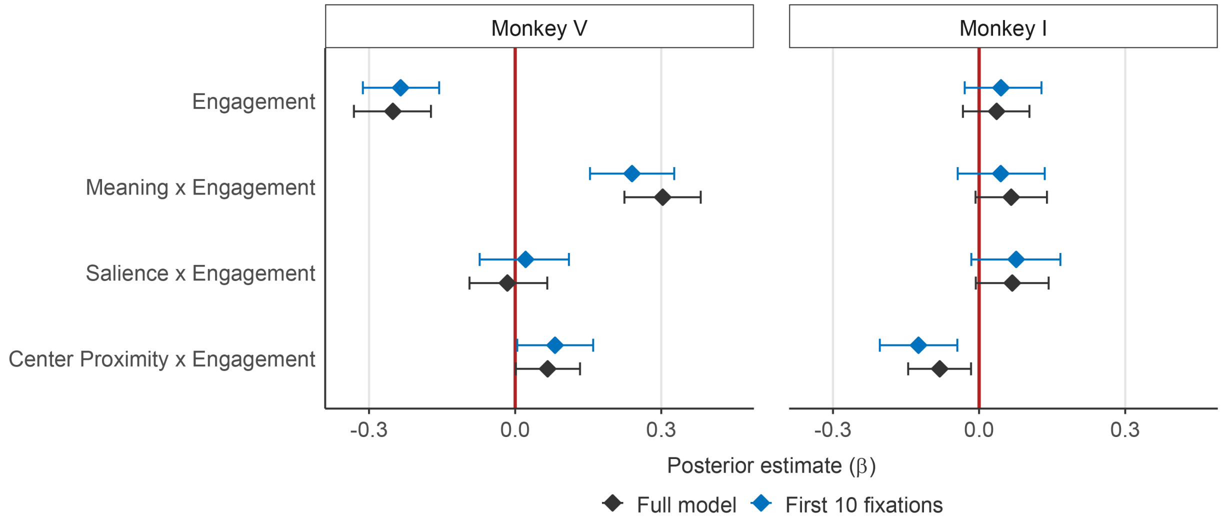


**Figure S12 | Engagement effects in the first 10 fixations of each trial.**

Posterior estimates of the engagement main effect and its interactions with each spatial predictor, for Monkey V (left) and Monkey I (right). Diamonds represent the posterior mean and horizontal bars the 95% HDI, for the full Engagement model (black) and for the model refitted using only the first 10 fixations of each scene (blue), which holds fixation count constant across trials. The vertical red line marks 0.

### Tables

| | | Monkey V | | | Monkey I | | |
|---|---|---|---|---|---|---|---|
| Metric | Model | Mean | SD | 95% HDI | Mean | SD | 95% HDI |
| *Bayes* $R^2$ | | | | | | | |
| $R^2$ | Fixation model | 0.26 | 0.01 | [0.25, 0.28] | 0.20 | 0.01 | [0.18, 0.21] |
| $R^2$ | Familiarity model | 0.27 | 0.01 | [0.25, 0.29] | 0.21 | 0.01 | [0.19, 0.23] |
| $R^2$ | Engagement model | 0.27 | 0.01 | [0.25, 0.28] | 0.20 | 0.01 | [0.18, 0.22] |
| *Posterior* $\mathrm{cor}(y, \mathrm{E}[y \mid x])$ | | | | | | | |
| $r$ | Fixation model | 0.48 | 0.0012 | [0.479, 0.484] | 0.43 | 0.0006 | [0.431, 0.433] |
| $r$ | Familiarity model | 0.50 | 0.0013 | [0.500, 0.505] | 0.46 | 0.0013 | [0.453, 0.458] |
| $r$ | Engagement model | 0.50 | 0.0010 | [0.499, 0.503] | 0.44 | 0.0009 | [0.433, 0.437] |

**Table S1 | Variance explained and predictive power for each model.**

The table summarizes model performance metrics for the Fixation model (Table 1), the Familiarity model (Table S2; fitted on the content-matched scenes), and the Engagement model (Table S5). The upper section reports the Bayesian $R^2$, representing the proportion of variance explained by the model's fixed effects. The lower section reports the posterior point-biserial correlation ($r$) between the observed binary fixation outcomes and the model's predicted probabilities. For each metric, the posterior mean, standard deviation (SD), and 95% highest density interval (HDI) are provided for Monkey V and Monkey I.

| | Monkey V | | | | Monkey I | | | |
|---|---|---|---|---|---|---|---|---|
| | $\beta$ Mean | $\beta$ SD | 95% HDI | $P(\beta > 0 \mid \mathrm{data})$ | $\beta$ Mean | $\beta$ SD | 95% HDI | $P(\beta > 0 \mid \mathrm{data})$ |
| *Fixed effects* | | | | | | | | |
| Intercept | -0.04 | 0.07 | [-0.17, 0.09] | 0.25 | 0.00 | 0.06 | [-0.12, 0.11] | 0.46 |
| Meaning | 0.97 | 0.08 | [0.82, 1.13] | $> 0.999$ | 0.85 | 0.07 | [0.71, 1.00] | $> 0.999$ |
| Salience | 0.67 | 0.07 | [0.54, 0.80] | $> 0.999$ | 0.49 | 0.06 | [0.37, 0.62] | $> 0.999$ |
| Center proximity | 0.06 | 0.06 | [-0.06, 0.17] | 0.84 | 0.33 | 0.06 | [0.21, 0.44] | $> 0.999$ |
| Familiarity | 0.09 | 0.09 | [-0.10, 0.27] | 0.83 | 0.04 | 0.08 | [-0.12, 0.19] | 0.67 |
| Meaning $\times$ salience | -0.20 | 0.09 | [-0.38, -0.03] | 0.01 | -0.25 | 0.07 | [-0.39, -0.11] | $< 0.001$ |
| Meaning $\times$ center proximity | 0.14 | 0.07 | [0.00, 0.28] | 0.97 | 0.01 | 0.07 | [-0.13, 0.16] | 0.56 |
| Salience $\times$ center proximity | 0.27 | 0.07 | [0.13, 0.42] | $> 0.999$ | 0.00 | 0.06 | [-0.12, 0.13] | 0.53 |
| Meaning $\times$ Familiarity | -0.26 | 0.11 | [-0.47, -0.06] | 0.01 | -0.47 | 0.10 | [-0.66, -0.28] | $< 0.001$ |
| Salience $\times$ Familiarity | 0.00 | 0.10 | [-0.19, 0.18] | 0.49 | 0.09 | 0.09 | [-0.08, 0.26] | 0.84 |
| Center proximity $\times$ Familiarity | 0.44 | 0.08 | [0.27, 0.60] | $> 0.999$ | 0.16 | 0.08 | [0.00, 0.32] | 0.97 |
| *Random effects* | | | | | | | | |
| Scene sigma | 0.23 | 0.08 | [0.05, 0.38] | $> 0.999$ | 0.10 | 0.07 | [0.00, 0.22] | $> 0.999$ |

**Table S2 | Posterior estimates for the Familiarity model.**

The fixed effects (main effects and two-way interactions) and the scene-level random effect are presented for the Fixation model (Table 1) extended to include scene familiarity, fitted on the content-matched scenes (standardized difference below 0.1; Figure S9 and Figure 4). Columns display the posterior mean and standard deviation (SD) of $\beta$ coefficients, the 95% highest density interval (HDI), and the probability of a positive effect ($P(\beta > 0 \mid \mathrm{data})$). Probabilities close to 1 indicate strong evidence for a positive effect, whereas probabilities close to 0 indicate strong evidence for a negative effect. Values above 0.999 or below 0.001 are reported as $> 0.999$ or $< 0.001$ to avoid implying exact certainty.

| | Monkey V | | | | Monkey I | | | |
|---|---|---|---|---|---|---|---|---|
| | All scenes | | Matched | | All scenes | | Matched | |
| Statistic | Std. diff. | $p$ | Std. diff. | $p$ | Std. diff. | $p$ | Std. diff. | $p$ |
| *Statistics used for matching* | | | | | | | | |
| Average meaning | -0.87 | $< 0.001$ | -0.08 | 0.774 | -0.85 | $< 0.001$ | -0.08 | 0.983 |
| Meaning SD (within scene) | -0.64 | $< 0.001$ | -0.05 | 0.595 | -0.53 | 0.001 | 0.00 | 0.842 |
| Salience SD (within scene) | 0.03 | 0.739 | 0.02 | 0.751 | 0.08 | 0.494 | -0.01 | 0.718 |
| Visual clutter | -0.57 | $< 0.001$ | -0.01 | 0.651 | -0.52 | 0.002 | -0.01 | 0.539 |
| Meaning–salience correlation | 0.18 | 0.227 | 0.03 | 0.844 | 0.29 | 0.060 | 0.02 | 0.851 |
| *Image statistics not used for matching* | | | | | | | | |
| Mean relative luminance | 0.14 | 0.449 | 0.26 | 0.193 | 0.16 | 0.288 | 0.31 | 0.115 |
| RMS contrast | -0.30 | 0.029 | -0.09 | 0.534 | -0.19 | 0.168 | 0.05 | 0.927 |
| Edge energy | -0.44 | 0.004 | 0.14 | 0.080 | -0.38 | 0.025 | 0.18 | 0.044 |
| Spectral slope | -0.13 | 0.393 | 0.27 | 0.070 | -0.10 | 0.523 | 0.27 | 0.068 |
| Colorfulness | -0.69 | $< 0.001$ | -0.66 | $< 0.001$ | -0.68 | $< 0.001$ | -0.70 | $< 0.001$ |
| Scenes (familiar / unfamiliar) | 95 / 98 | | 62 / 62 | | 82 / 93 | | 59 / 59 | |

**Table S3 | Comparison of familiar and unfamiliar scenes on ten scene statistics.**

Familiar and unfamiliar scenes are compared on the five statistics used for matching (the average meaning of the scene, the within-scene standard deviation of meaning and of salience, visual clutter, and the within-scene correlation between meaning and salience) and on five low-level image statistics that were not used for matching (mean relative luminance, RMS contrast, edge energy, spectral slope, and colorfulness). Each comparison is reported for all analyzed scenes and for the matched set of each animal, as a standardized difference and as a Wilcoxon rank-sum test. The standardized difference is the familiar minus the unfamiliar mean, divided by the standard deviation across all analyzed scenes. Colorfulness was the only statistic that still differed reliably between conditions in both animals after matching, and edge energy still differed in Monkey I. Each was therefore added to the Familiarity model as a covariate in a separate fit (Table S4).

| | Monkey V | | | | Monkey I | | | |
|---|---|---|---|---|---|---|---|---|
| | $\beta$ Mean | $\beta$ SD | 95% HDI | $P(\beta > 0 \mid \text{data})$ | $\beta$ Mean | $\beta$ SD | 95% HDI | $P(\beta > 0 \mid \text{data})$ |
| **Familiarity model with colorfulness** | | | | | | | | |
| *Familiarity terms* | | | | | | | | |
| Familiarity | 0.08 | 0.10 | [-0.11, 0.27] | 0.79 | 0.00 | 0.09 | [-0.16, 0.18] | 0.50 |
| Meaning × Familiarity | -0.32 | 0.11 | [-0.54, -0.09] | 0.003 | -0.58 | 0.10 | [-0.78, -0.38] | $< 0.001$ |
| Salience × Familiarity | -0.05 | 0.10 | [-0.25, 0.15] | 0.31 | 0.08 | 0.09 | [-0.10, 0.26] | 0.81 |
| Center proximity × Familiarity | 0.50 | 0.09 | [0.33, 0.67] | $> 0.999$ | 0.16 | 0.09 | [-0.01, 0.34] | 0.97 |
| *Colorfulness terms* | | | | | | | | |
| Colorfulness | -0.01 | 0.04 | [-0.10, 0.08] | 0.42 | -0.05 | 0.04 | [-0.13, 0.03] | 0.10 |
| Meaning × colorfulness | -0.12 | 0.05 | [-0.23, -0.02] | 0.01 | -0.21 | 0.05 | [-0.30, -0.11] | $< 0.001$ |
| Salience × colorfulness | -0.07 | 0.05 | [-0.16, 0.02] | 0.07 | -0.01 | 0.05 | [-0.10, 0.08] | 0.40 |
| Center proximity × colorfulness | 0.10 | 0.04 | [0.02, 0.18] | 0.99 | 0.04 | 0.04 | [-0.04, 0.12] | 0.82 |
| **Familiarity model with edge energy** | | | | | | | | |
| *Familiarity terms* | | | | | | | | |
| Familiarity | 0.15 | 0.09 | [-0.02, 0.32] | 0.96 | 0.07 | 0.08 | [-0.08, 0.23] | 0.82 |
| Meaning × Familiarity | -0.32 | 0.10 | [-0.52, -0.11] | 0.001 | -0.52 | 0.10 | [-0.71, -0.33] | $< 0.001$ |
| Salience × Familiarity | 0.04 | 0.10 | [-0.15, 0.23] | 0.67 | 0.11 | 0.09 | [-0.06, 0.29] | 0.90 |
| Center proximity × Familiarity | 0.46 | 0.08 | [0.30, 0.63] | $> 0.999$ | 0.20 | 0.08 | [0.03, 0.36] | 0.99 |
| *Edge energy terms* | | | | | | | | |
| Edge energy | -0.29 | 0.05 | [-0.38, -0.19] | $< 0.001$ | -0.17 | 0.04 | [-0.24, -0.09] | $< 0.001$ |
| Meaning × edge energy | 0.15 | 0.05 | [0.04, 0.25] | 0.998 | -0.10 | 0.05 | [-0.19, -0.01] | 0.01 |
| Salience × edge energy | -0.13 | 0.05 | [-0.22, -0.04] | 0.003 | -0.04 | 0.05 | [-0.12, 0.05] | 0.21 |
| Center proximity × edge energy | 0.00 | 0.04 | [-0.08, 0.08] | 0.54 | -0.02 | 0.04 | [-0.11, 0.05] | 0.28 |

**Table S4 | Posterior estimates for the Familiarity model with colorfulness or edge energy as a covariate.**

The table shows two models. The first is the Familiarity model fitted on the matched scenes with colorfulness added. The second is the same model with edge energy added instead. In each model, the added statistic was $z$-scored across the sampled locations and included as a main effect and in interaction with meaning, salience, and center proximity. For each model, the familiarity terms and the terms of the added statistic are shown. Columns display the posterior mean and standard deviation (SD) of $\beta$ coefficients, the 95% highest density interval (HDI), and the probability of a positive effect ($P(\beta > 0 \mid \text{data})$). Values above 0.999 or below 0.001 are reported as $> 0.999$ or $< 0.001$ to avoid implying exact certainty. The meaning $\times$ familiarity interaction observed in the Familiarity model (Table S2) was retained in both animals in both models.

| | Monkey V | | | | Monkey I | | | |
|---|---|---|---|---|---|---|---|---|
| | $\beta$ Mean | $\beta$ SD | 95% HDI | $P(\beta > 0 \mid \mathrm{data})$ | $\beta$ Mean | $\beta$ SD | 95% HDI | $P(\beta > 0 \mid \mathrm{data})$ |
| *Fixed effects* | | | | | | | | |
| Intercept | -0.05 | 0.04 | [-0.13, 0.02] | 0.08 | 0.05 | 0.04 | [-0.01, 0.12] | 0.94 |
| Meaning | 0.72 | 0.05 | [0.63, 0.82] | $> 0.999$ | 0.47 | 0.04 | [0.38, 0.55] | $> 0.999$ |
| Salience | 0.69 | 0.04 | [0.62, 0.77] | $> 0.999$ | 0.60 | 0.04 | [0.52, 0.67] | $> 0.999$ |
| Center proximity | 0.31 | 0.03 | [0.24, 0.38] | $> 0.999$ | 0.39 | 0.03 | [0.32, 0.46] | $> 0.999$ |
| Engagement | -0.25 | 0.04 | [-0.33, -0.17] | $< 0.001$ | 0.04 | 0.04 | [-0.03, 0.10] | 0.85 |
| Meaning $\times$ salience | -0.22 | 0.04 | [-0.31, -0.14] | $< 0.001$ | -0.20 | 0.04 | [-0.27, -0.13] | $< 0.001$ |
| Meaning $\times$ center proximity | 0.07 | 0.04 | [0.00, 0.15] | 0.97 | 0.00 | 0.04 | [-0.07, 0.06] | 0.45 |
| Salience $\times$ center proximity | 0.08 | 0.04 | [0.00, 0.16] | 0.98 | 0.04 | 0.04 | [-0.03, 0.11] | 0.86 |
| Meaning $\times$ engagement | 0.30 | 0.04 | [0.22, 0.38] | $> 0.999$ | 0.07 | 0.04 | [-0.01, 0.14] | 0.96 |
| Salience $\times$ engagement | -0.02 | 0.04 | [-0.09, 0.07] | 0.35 | 0.07 | 0.04 | [-0.01, 0.14] | 0.96 |
| Center proximity $\times$ engagement | 0.07 | 0.03 | [0.00, 0.13] | 0.98 | -0.08 | 0.03 | [-0.15, -0.02] | 0.01 |
| *Random effects* | | | | | | | | |
| Scene sigma | 0.30 | 0.06 | [0.19, 0.41] | $> 0.999$ | 0.18 | 0.08 | [0.01, 0.30] | $> 0.999$ |

#### Table S5 | Posterior estimates for the Engagement model.

The fixed effects (main effects and two-way interactions) and the scene-level random effect are presented for the Fixation model (Table 1) extended to include scene engagement (total fixation duration per trial). Columns display the posterior mean and standard deviation (SD) of the $\beta$ coefficients, the 95% highest density interval (HDI), and the probability of a positive effect ($P(\beta > 0 \mid \mathrm{data})$). Probabilities close to 1 indicate strong evidence for a positive effect, whereas probabilities close to 0 indicate strong evidence for a negative effect. Values above 0.999 or below 0.001 are reported as $> 0.999$ or $< 0.001$ to avoid implying exact certainty.

| | | Monkey V | Monkey I |
|---|---|---|---|
| Model | Parameter pair | $r$ | $r$ |
| *Fixation model* | | | |
| | Meaning vs. Salience | -0.16 | -0.13 |
| | Meaning vs. Center proximity | -0.29 | -0.24 |
| | Salience vs. Center proximity | 0.10 | 0.05 |
| *Familiarity model* | | | |
| | Meaning vs. Salience | -0.12 | -0.21 |
| | Meaning vs. Center proximity | -0.39 | -0.27 |
| | Meaning vs. Familiarity | 0.08 | 0.08 |
| | Salience vs. Center proximity | 0.07 | 0.06 |
| | Salience vs. Familiarity | 0.03 | 0.04 |
| | Center proximity vs. Familiarity | -0.06 | -0.08 |
| *Engagement model* | | | |
| | Meaning vs. Salience | -0.12 | -0.11 |
| | Meaning vs. Center proximity | -0.30 | -0.23 |
| | Meaning vs. Engagement | -0.25 | -0.09 |
| | Salience vs. Center proximity | 0.06 | 0.06 |
| | Salience vs. Engagement | -0.03 | 0.02 |
| | Center proximity vs. Engagement | 0.05 | 0.07 |

**Table S6 | Posterior parameter correlations.**

Pairwise correlations ($r$) between the posterior estimates of fixed-effect predictors across the Fixation, Familiarity, and Engagement models. Values are modest ($|r| < 0.40$) for both subjects, indicating that the model separates the effects of the predictors well. These are correlations between the parameter estimates, not between the predictors themselves (Figure S2).

| | | Monkey V | | Monkey I | |
|---|---|---|---|---|---|
| Model | Term | ESS (Bulk) | $\hat{R}$ | ESS (Bulk) | $\hat{R}$ |
| *Fixation model* | | | | | |
| | Meaning | 8,989 | 1.00 | 6,428 | 1.00 |
| | Salience | 21,625 | 1.00 | 18,018 | 1.00 |
| | Center proximity | 20,443 | 1.00 | 24,432 | 1.00 |
| *Temporal model* | | | | | |
| | Meaning | 6,007 | 1.00 | 5,720 | 1.00 |
| | Salience | 5,597 | 1.00 | 5,965 | 1.00 |
| | Center proximity | 6,199 | 1.00 | 5,826 | 1.00 |
| | Fixations 4–6 | 8,534 | 1.00 | 11,031 | 1.00 |
| | Fixations 7–10 | 7,965 | 1.00 | 10,916 | 1.00 |
| | Fixations 11 or later | 7,075 | 1.00 | 10,229 | 1.00 |
| *Familiarity model* | | | | | |
| | Meaning | 12,698 | 1.00 | 16,746 | 1.00 |
| | Salience | 16,661 | 1.00 | 18,034 | 1.00 |
| | Center proximity | 14,819 | 1.00 | 18,607 | 1.00 |
| | Familiarity | 17,411 | 1.00 | 27,454 | 1.00 |
| *Engagement model* | | | | | |
| | Meaning | 9,251 | 1.00 | 6,612 | 1.00 |
| | Salience | 19,159 | 1.00 | 17,116 | 1.00 |
| | Center proximity | 17,512 | 1.00 | 27,115 | 1.00 |
| | Engagement | 13,809 | 1.00 | 21,934 | 1.00 |

**Table S7 | Markov chain Monte Carlo (MCMC) sampling diagnostics.**

Effective Sample Size (ESS, bulk) and Gelman–Rubin convergence statistics ($\hat{R}$) for the main predictors in the Fixation, Temporal, Familiarity, and Engagement models. All $\hat{R}$ values are 1.00 and ESS values exceed 5500, confirming stable convergence of the MCMC chains.
